# Attention affects overall gain but not selective contrast at meter frequencies in the neural processing of rhythm

**DOI:** 10.1101/2020.09.23.309443

**Authors:** Tomas Lenc, Peter E. Keller, Manuel Varlet, Sylvie Nozaradan

## Abstract

When listening to music, humans spontaneously perceive and synchronize movement to periodic pulses of meter. A growing body of evidence suggests that this widespread ability is related to neural processes that selectively enhance meter periodicities. However, to what extent these neural processes are affected by the attentional state of the listener remains largely unknown. Here, we recorded EEG while participants listened to auditory rhythms and detected small changes in tempo or pitch of the stimulus, or performed a visual task. The overall neural response to the auditory input decreased when participants attended the visual modality, indicating generally lower sensitivity to acoustic information. However, the selective contrast at meter periodicities did not differ across the three tasks. Moreover, this selective contrast could be trivially accounted for by biologically-plausible models of subcortical auditory processing, but only when meter periodicities were already prominent in the acoustic input. However, when meter periodicities were not prominent in the auditory input, the EEG responses could not be explained by low-level processing. This was also confirmed by early auditory responses that originate predominantly in early auditory areas and were recorded in the same EEG. The contrast at meter periodicities in these early responses was consistently smaller than in the EEG responses originating mainly from higher-level processing stages. Together, these results demonstrate that selective contrast at meter periodicities involves higher-level neural processes that may be engaged automatically, irrespective of behavioral context. This robust shaping of the neural representation of rhythm might thus contribute to spontaneous and effortless synchronization to musical meter in humans across cultures.

## Introduction

Perception of rhythmic sound sequences involves much more than just a precise representation of constituent time intervals. Already perception of single intervals is not one-to-one with respect to the sensory input, but reflects a representation constructed with respect to prior individual experience (Desain and Honing, 2003; Jazayeri and Shadlen, 2010; Jacoby and McDermott, 2017). An even higher level of perceptual organization is arguably at stake when the rhythmic input induces perception of musical meter, i.e., a nested set of periodic pulses to which people tend to move or dance (Cohn, 2020). That is, the internal representation of meter guides perceptual organization of the incoming rhythmic sequence in time (Povel and Essens, 1985; McAuley and Jones, 2003) and drives body movement such as head bobbing or foot tapping (Toiviainen et al., 2010; Janata et al., 2012). Perception and sensory-motor synchronization to meter is a spontaneous human ability that has been widely observed across cultures and musical traditions (Nettl, 2000; Savage et al., 2015).

In some cases, meter perception can be largely driven by the acoustic features of the sensory input, particularly when clear periodicities are present in the temporal structure of the stimulus (although even in such cases the alignment of the perceived pulses with the input is not trivial, see e.g. off-beat rhythm in reggae). However, meter perception is often induced by stimuli that lack unambiguous acoustic cues to meter periodicities (Chapin et al., 2010; Nozaradan et al., 2012; Witek et al., 2014; Large et al., 2015; London et al., 2017; Vuust et al., 2018; Matthews et al., 2020), and the same rhythmic sequence can be perceptually organized in different ways depending on prior experience at multiple timescales (Phillips-Silver and Trainor, 2005; Hannon et al., 2012a; Chemin et al., 2014; van der Weij et al., 2017). This shows that meter perception goes beyond the mere tracking of periodicities in the sensory input, and additionally involves higher-level processes that transform the input towards a particular metric category with a great degree of robustness and flexibility with respect to the input (Nozaradan et al., 2017a).

This is in line with a number of recent neurophysiological studies based on the assumption that meter perception is related to neural processes that emphasize the contrast between time points marked by the perceived metric pulses and other time points not marked by pulses. This contrast in the neural response can be driven already by the physical features of the sensory input along with a set of low-level nonlinear transformations throughout early auditory processing stages (Rajendran et al., 2017, 2020). Importantly there is also increasing evidence for higher-level neural processes that transform the input by selectively enhancing this contrast beyond physical features and low-level nonlinearities (Lenc et al., 2018, 2020). These higher-level neural processes may thus play a key role in building internal representation of meter dissociated from the physical features of the sensory input (Nozaradan et al., 2011, 2012, 2017a, 2017b; Tal et al., 2017).

However, to what extent these processes are engaged automatically, and whether they depend on the behavioral goals of the listener remains largely unknown. Previous neurophysiological and neuroimaging studies of meter processing in humans have employed a wide range of behavioral tasks, some instructing participants to attend directly to the pulse-like metric structure of the stimuli (Grahn and Rowe, 2009, 2013; Lenc et al., 2020; Matthews et al., 2020) or the temporal properties of the stimulus (Nozaradan et al., 2017b; Lenc et al., 2018), while other studies used an orthogonal task such as attending to a non-temporal sound feature (e.g. pitch; Haumann et al., 2018) or attending to a different modality (e.g. visual; Chapin et al., 2010) or no task at all (Bengtsson et al., 2009). However, how neural processing of a rhythmic input changes across these different tasks has not been systematically explored using a consistent set of stimuli and analysis methods.

Additionally, in a series of studies investigating putative “pre-attentive beat perception” using event-related brain response to regularity violations, participants were typically asked to perform a passive task, such as watching a silent movie, while listening to the rhythmic stimuli (Vuust et al., 2005; Ladinig et al., 2009; Geiser et al., 2010; Bouwer et al., 2014, 2016). The lack of strict control of participant’s attentional focus combined with the low load of the task make the results of these studies difficult to interpret (Lavie and Dalton, 2014; Sussman et al., 2014; Murphy et al., 2017). In addition, these studies mostly used stimuli with clear acoustic cues to meter periodicities. Therefore, it remains unknown whether these results would generalize to rhythmic inputs that lack such prominent sensory cues and may thus require higher-level processes to induce meter perception (Chapin et al., 2010; Nozaradan et al., 2011).

In the current study, we aimed to address these issues by recording human brain electroencephalographic (EEG) activity in response to (i) a consistent set of rhythmic stimuli with varying amounts of sensory cues to meter periodicities, along with (ii) a set of three demanding behavioral tasks in the same sample of participants. We presented participants with two rhythmic sequences. One sequence contained prominent acoustic cues to meter periodicities, while the other sequence lacked such prominent periodic cues. This latter sequence enabled us to control for a low-level confound which could trivially explain enhanced neural response at meter periodicities. That is, if selective contrast at meter periodicities is observed in the EEG in response to a sequence lacking such prominent periodic cues, this selective contrast at meter periodicities cannot be explained easily by the stimulus structure or low-level processing of the stimulus. Importantly, the decision as to what frequencies would correspond to meter periodicities was informed by previous studies, which used tapping tasks to carefully test the metric pulses most consistently induced by these two rhythmic patterns across listeners (Nozaradan et al., 2012, 2018; Lenc et al., 2018). This ensured that these specific frequencies were relevant for meter perception, in contrast to other frequencies that are also elicited by the rhythms but are irrelevant to the perceived meter.

Participants listened to the rhythms while performing three different demanding tasks. In the first task, participants were required to detect small changes in the speed of the rhythmic sequence. Because the sequence was non-isochronous, this task cannot be carried out by simply comparing successive inter-tone intervals and therefore encourages participants to build an internal representation of meter that aids tracking of the overall speed of the rhythm (Schulze, 1978; Grube and Griffiths, 2009; Grube et al., 2010). In the second task, participants were required to detect small changes in the pitch of a single tone among the rhythmic sequences, thus still focusing on the sound but not necessarily on its timing. Finally, in the third task, participants were required to mentally sum numbers sequentially presented on the screen while ignoring the sounds altogether.

The EEG was recorded while participants were presented with the auditory sequences and carried out the behavioral tasks without any movement. From the EEG, we measured the difference in amplitude of the neural activity at meter-related frequencies vs. meter-unrelated frequencies elicited by the rhythms, i.e., the contrast at perceptually-relevant timescales, using frequency tagging. This approach has proven to be a powerful tool for capturing the contrast in brain responses between periodically spaced time points with high signal-to-noise ratio and without assumptions about the latency or the shape of the response (Nozaradan, 2014; Rossion, 2014; Norcia et al., 2015; Nozaradan et al., 2017a; Rossion et al., 2020). Moreover, the approach also allows the overall gain of the response (i.e. the general sensitivity to auditory stimulation) to be disentangled from the selective contrast at meter-relevant periodicities.

We also examined whether the contrast at meter frequencies in the EEG activity elicited across behavioral tasks could be trivially accounted for by fixed nonlinear transformations along the early auditory pathway. To this end, we used biologically plausible models to simulate responses to the rhythmic stimuli in the auditory nerve, as well as inferior colliculus. To complement these simulations, we also directly captured responses presumed to be predominantly driven by brainstem auditory nuclei and primary auditory areas using the same frequency-tagging method as Nozaradan et al. (2016c, 2018). These early responses were observed at a faster timescale (> 150 Hz) due to neural tracking of the amplitude-modulated fine structure of the sound input. By comparing these early auditory responses to the higher-level responses observed at slower timescales (< 5Hz, corresponding to the amplitude envelope of the input), which mainly capture activity in higher-level cortical networks, we aimed to estimate the contribution of different processing stages to the selective contrast at meter frequencies across different attentional contexts.

## Materials and methods

### Participants

Seventeen healthy volunteers (mean age = 23.3, SD = 6.7, 15 females) with various levels of formal musical training (mean = 2.5, SD = 4.6, range = 0-16 years) participated in the study after providing written informed consent. All participants reported normal hearing and no history of neurological or psychiatric disorder. The study was approved by the Research Ethics Committee of Western Sydney University.

### Auditory stimuli

The auditory stimuli were created in Matlab R2016b (The MathWorks, Natick, MA) and presented binaurally through insert earphones (ER-2; Etymotic Research, Elk Grove Village, IL) at a comfortable listening level (∼ 75 dB SPL) using PsychToolbox, version 3.0.14 (Brainard, 1997) running on a MacBook Pro laptop (mid-2015, OSX 10.12). Triggers were sent to the EEG system using LabJack U3 interface. The stimuli consisted of a 2.4-s long rhythmic pattern (made up of twelve 200-ms long events) continuously looped 14 times to create a 33.6-s long sequence. The rhythmic structure of the pattern was based on a specific arrangement of 8 sound events and 4 silent events (amplitude at 0). Each sound event corresponded to a complex tone consisting of three partials (f1 = 209 Hz, f2 = 398 Hz, f3 = 566 Hz) with linear onset and offset ramp lasting 10% of the event duration (i.e. 20 ms).

We used two different rhythmic patterns (depicted in Figure 1). These two patterns were selected based on previous evidence that they both induce a perception of musical meter, consistent across individuals, based on nested grouping of the individual event rate (200 ms) by 2 (2 x 200 ms = 400 ms), 2 (2 x 400 ms = 800 ms) and 3 (3 x 800 ms = 2400 ms) (Nozaradan et al., 2012, 2018; Lenc et al., 2018).

**Figure 1.**
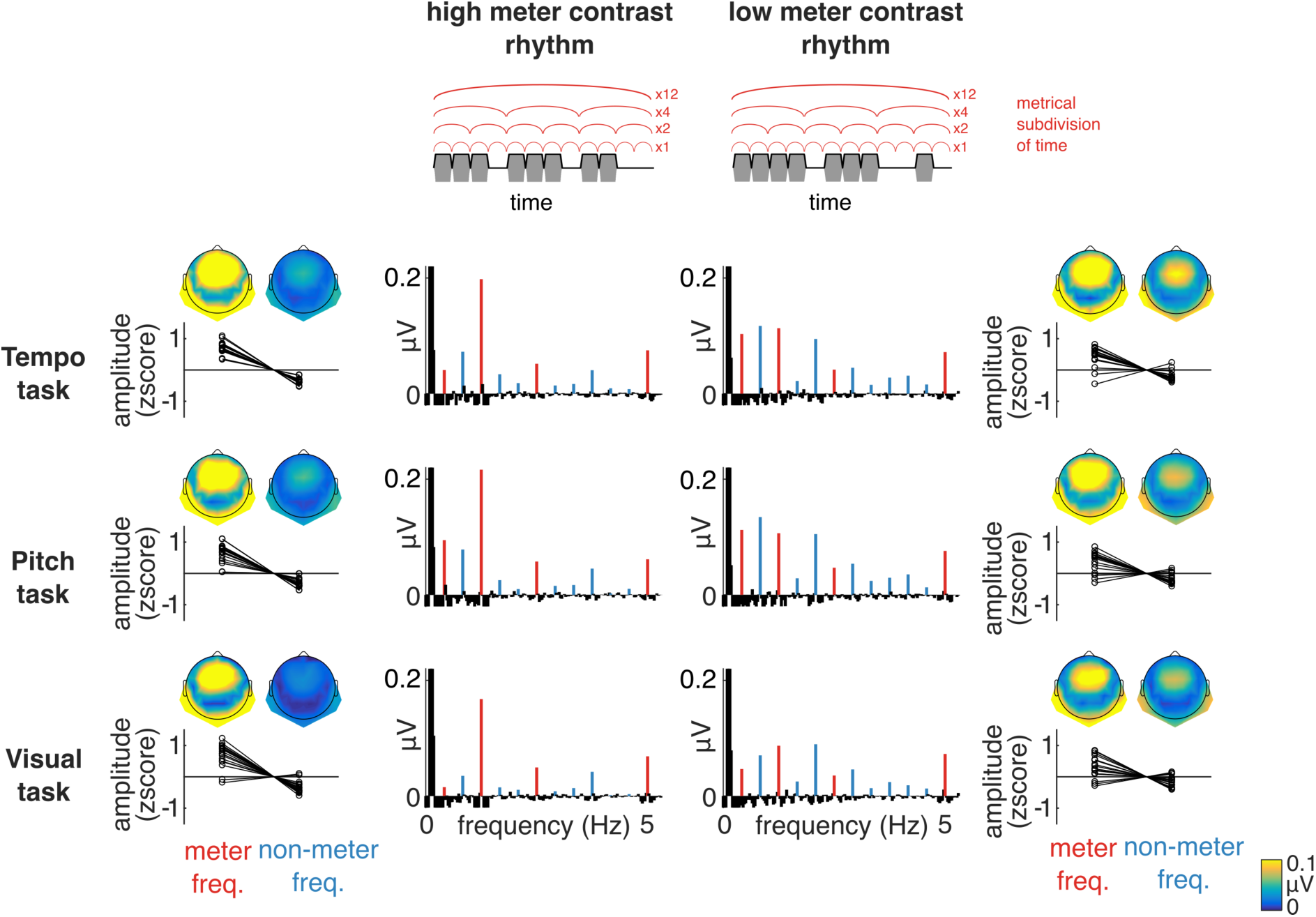
Stimulus design and higher-level EEG responses. (Top) The sound waveform representing one cycle of the high meter contrast (Left) and low meter contrast (Right) rhythmic pattern is depicted in grey. The broadband envelope is overlaid as a black line. Above each pattern, the meter typically induced by these patterns is shown as red arches representing individual pulses in the metric structure. (Bottom) Spectra of higher-level EEG responses elicited for each rhythm and task (average across all participants and EEG channels). Mean z-scored amplitude elicited at meter-related (red) and meter-unrelated (blue) frequencies is shown next to the corresponding spectra (data points represent individual participants), along with the topographical distribution of mean EEG amplitude at these two subsets of frequencies (average across all participants).

Importantly, although the two rhythmic patterns induce perception of musical meter at consistent periods across individuals, they provide the listener with different amounts of direct sensory cues to this perceived metric structure. One way to quantify this is to examine the degree of mismatch between the perceived meter and the arrangement of sound events in the rhythm using syncopation scores. Even though different ways to calculate syncopation scores have been proposed, the main principle they share is quantifying to what extent the rhythmic stimulus creates a contrast between time points that coincide with the putative metric pulses, and the rest of the time points, i.e. a contrast at meter periodicities (Longuet-Higgins and Lee, 1984; Povel and Essens, 1985; Parncutt, 1994; Eck, 2003). We calculated syncopation scores for the two rhythmic patterns using an algorithm originally proposed by Longuet-Higgins and Lee, which simultaneously takes into account the whole nested hierarchy of metric pulses (Longuet-Higgins and Lee, 1984; Witek et al., 2014). Additionally, a C score (counterevidence) was calculated using the method and parameters proposed by Povel and Essens (1985). While C score calculates syncopation using only one pulse in the metric structure, it accounts for variable perceptual salience of tones making up the pattern based on their relative temporal proximity (Povel and Okkerman, 1981).

Even though the periods of the perceived metric pulses for the two rhythms are generally consistent across participants, the alignment of these pulses with respect to the rhythmic stimulus can vary (Nozaradan et al., 2012, 2018; Lenc et al., 2018). To avoid assumptions regarding particular pulse alignment, the minimum syncopation and C score across all 12 possible positions of the slowest metric pulse with respect to the rhythm was taken (Lenc et al., 2020). This yielded smaller scores for one rhythm (syncopation = 1, C = 1), in comparison to the other rhythm (syncopation = 2, C = 2). In other words, both measures revealed a greater mismatch between the perceived meter and the arrangement of sound events for the second rhythm.

This reflects the fact that the physical structure of the first rhythm provides clear and unambiguous information about the perceived meter. On the other hand, the second rhythm provides less sensory information about the metric periodicities (there is no plausible alignment of the perceived pulses that would lead to systematic match with the distribution of sound onsets in the pattern). For these reasons, the first and the second rhythm are further referred to as “high meter contrast” and “low meter contrast” rhythm, respectively (note that various terms have been previously used to describe these same rhythms, e.g. unsyncopated and syncopated, Nozaradan et al., 2016b, 2017b, 2018). Despite these differences, both rhythms consistently induce meter perception across listeners, as revealed by previous studies (Nozaradan et al., 2012, 2018; Lenc et al., 2018).

### Frequency-tagging analysis

Another way to measure the amount of contrast at meter periodicities is to directly analyze the modulation spectrum of the acoustic stimulus using Fourier transform. This allows quantification of the extent to which the continuous modulation of acoustic features of the input (here amplitude envelope) emphasizes particular periodicities.

Because the stimulus sequence consisted of seamless repetitions of the same rhythmic pattern, the modulation spectra were expected to contain energy at frequencies corresponding to the repetition of the pattern (1/2.4 s = 0.416 Hz) and harmonics. The relative distribution of energy across these different harmonics reveals how much contrast was present in the signal modulations at the corresponding frequencies. From the set of first 12 harmonics (up to 5 Hz, the frequency of individual event rate in the rhythms), four frequencies were considered meter-related (0.416, 1.25, 2.5, 5 Hz), as they corresponded to the frequencies of the perceived metric pulses (1/2.4 s, 1/0.8 s, 1/0.4 s, 1/0.2 s respectively). The remaining 8 frequencies in the set were considered meter-unrelated.

To measure the relative prominence of meter frequencies, amplitudes at the 12 frequencies corresponding to the stimulus modulation spectrum were converted to z-scores as follows: ([x] − [mean across the 12 frequencies])/[SD across the 12 frequencies]. A higher z-score at a specific frequency indicates that the response at that frequency stands out prominently relative to the whole set of frequencies in the modulation spectrum. The z-scores for meter-related frequencies were averaged to obtain an index of their relative prominence in the modulation spectra.

The main advantage of using FFT is that it can be applied to a variety of signals representing (i) modulations in the acoustic input, (ii) simulated responses of neurons in the subcortical auditory nuclei, (iii) surface EEG, and (iv) movement. Importantly, using the z-scoring standardization yields a measure invariant to differences in unit and scale, thus allowing for objective measurement of the relative distance between these different signals. In sum, this method represents a powerful tool to track the transformation of the input, i.e. the changes in contrast at meter periodicities across different processing stages from input to output.

### Models of subcortical auditory processing

To estimate to what extent the neural transformation of a rhythmic acoustic stimulus could be driven by early stages of the auditory pathway, we simulated responses to the rhythmic stimuli using multiple biologically-plausible models of subcortical auditory processing, as described below. Comparing the EEG responses to these early representations thus helps to disentangle the contribution of higher-level transformations that cannot be trivially explained by early sound processing stages.

#### (i) Broadband envelope

A number of previous EEG studies used broadband envelopes to represent modulations in the acoustic input (Aiken and Picton, 2008; Nozaradan et al., 2012, 2018; Chemin et al., 2014; Cirelli et al., 2016; Tal et al., 2017; Broderick et al., 2019; Di Liberto et al., 2020). To provide a point of comparison with these studies, the broadband amplitude envelope of the 33.6-s auditory sequences (high and low meter contrast rhythm) was extracted using the Hilbert transform (as implemented in Matlab) and then transformed into the frequency domain using a fast Fourier transform (FFT, yielding a spectral resolution of 1/33.6 s, i.e. approximately 0.03 Hz).

#### (iia) UR-EAR-AN

The model of the auditory nerve developed by Bruce et al. (2018) as implemented in UR_EAR toolbox (version 2020a) was used to simulate responses from 128 cochlear channels with characteristic frequencies logarithmically spaced between 130 and 8000 Hz. The parameters used for cochlear tuning matched data available from human subjects (Shera et al., 2002). For each channel, 51 auditory nerve fibers were simulated with biologically plausible distribution of high, mid, and low-spontaneous-rate fibers (Liberman, 1978). The model provides faithful simulation of physiological processes associated with cochlear nonlinearities, inner hair cell transduction process, the synapse between the hair cell and the auditory nerve, and the associated firing rate adaptation.

#### (iib) UR-EAR-IC

The simulated auditory nerve firing rates were fed into the same-frequency inhibition and excitation model (SFIE) used to simulate enhanced onset synchrony and the decreased upper limit for phase-locking to stimulus envelope in the ventral cochlear nucleus (Nelson and Carney, 2004). The default parameters in the UR_EAR toolbox were used, which were based on Carney et al. (2015). A second SFIE model was then used to simulate band-pass modulation filtering and enhanced onset responses of neurons in the inferior colliculus (IC). The parameters were set to simulate IC units with the best modulation frequencies separately at 2, 4, 8, 16, 32, and 64 Hz.

For both the AN and IC stage of the UR_EAR model, the simulated instantaneous firing rates were summed across cochlear channels (Zuk et al., 2018; Rajendran et al., 2020), and transformed into the frequency domain using FFT. While averaging firing rates across channels might yield different results than averaging FFT magnitudes for spectrally complex inputs, the two methods should give very similar results for the stimuli in the current study, as the modulation waveform was identical across the whole spectrum. Subsequently, amplitudes at the 12 frequencies of interest were extracted from the obtained spectra and normalized by z-scoring separately for each model output (see section Frequency-tagging analysis).

### Early auditory responses

The frequencies of the partials of the complex tones delivering the rhythm (f1 = 209 Hz, f2 = 398 Hz, f3 = 566 Hz) were selected because sustained frequency-following responses at these frequencies are expected to originate predominantly from sub-cortical auditory nuclei due to low-pass characteristics of the ascending auditory pathway (Chandrasekaran and Kraus, 2010; Skoe and Kraus, 2010; but see Coffey et al., 2016, 2019, who show that a portion of this response could also be explained by activity from early cortical stages). Non-harmonic spacing of the partials was used in the current study as it is expected to elicit responses at frequencies that are not physically present in the stimulus spectrum. These frequencies corresponded to distortion-product otoacoustic emissions generated by nonlinear processes at the cochlear level and transmitted along the ascending auditory pathway (Lee et al., 2009). Hence, any EEG response at these frequencies could not be explained by an electromagnetic artifact from the sound-delivery system. These responses were expected at frequencies corresponding to quadratic distortion products across the three partials, i.e. f2-f1 (168 Hz), f3-f2 (189 Hz), and f3-f1 (357 Hz). Due to the frequency-shifting theorem, each of the distortion-product frequencies was expected to be symmetrically flanked by sidebands representing the amplitude modulation spectrum of the response (Oppenheim and Schafer, 2009). This allowed the contrast at meter frequencies to be quantified at earlier auditory processing stages with the same method as described above for the sound input (see section Frequency-tagging analysis). Furthermore, this contrast at meter frequencies obtained from earlier auditory stages was also compared to the contrast at meter frequencies obtained from EEG responses measured in a much lower frequency range (here at 5 Hz and below) and assumed to predominantly originate from higher-level processing stages (further referred to as “higher-level” responses) (Nozaradan et al., 2018). Importantly, because the index of contrast at meter frequencies consists in a relative measure of the amplitude at meter frequencies vs. meter-unrelated frequencies obtained after z-scoring standardization, this measure is invariant to differences in unit and scale, thus providing valid estimation of the relative distance between signals as different as the early auditory responses and higher-level responses, irrespective of differences in overall gain.

### Experimental design and procedure

Participants were presented with the rhythmic auditory stimuli in separate blocks of 10 self-paced trials. The polarity of the acoustic waveform was inverted on every other trial to prevent potential electromagnetic artifact at the frequencies of the sound input (Skoe and Kraus, 2010). In each block, participants were asked to perform a specific behavioral task.

#### Tempo task

The block contained two additional randomly-placed trials where one rhythm cycle (at a random position after the first 3 cycles) contained a decrease in tempo. This was implemented by gradually increasing (and then decreasing) the inter-onset intervals of the individual constituent events within one rhythm cycle according to a cosine window from the standard inter-onset interval (200 ms) to the maximum interval determined individually for each participant. Participants were asked to focus on the tempo of the stimuli, while ignoring all other parameters, as well as any visually presented stimuli. They reported whether the change was present after the end of each trial.

#### Pitch task

The block contained two additional trials with increased pitch of a single constituent tone (implemented as a proportional increase in the frequency of each partial). Participants were asked to report the presence of the pitch change at the end of each trial, while ignoring other sound parameters and visual stimuli.

#### Visual stimuli and task

Throughout all trials and blocks, participants also viewed sequentially presented numbers in the center of the screen positioned in front of them (approximately one meter distance). The numbers were randomly sampled such that the first number for each trial was between 100 and 200, and all subsequent numbers were between 10 and 30. The time interval between the onset of each sequential number was individually determined for each participant, and a jitter of 10% of this time interval was then applied to avoid any strict periodicity in the visual presentation of the numbers, which could result in a narrow frequency peak elicited in the EEG spectrum at the frequency of the visual presentation. Each number stayed on the screen for 80% of the inter-onset interval (with 10% random jitter applied to this value). Participants were asked to fixate their eyes on the numbers in every trial across all blocks in order to prevent eye movements. During the Visual task, they were asked to mentally add these numbers and report the sum at the end of each trial, while ignoring the sound stimuli. Participants were instructed to keep adding the incoming numbers even in case they missed any. This was to make sure participants did not “give up” in the middle of the trial, but kept continuously engaged with the visual task.

Each task and rhythm were presented as a separate block, yielding 3 x 2 = 6 blocks in the whole EEG session (block order was counterbalanced across participants). Participants were seated in a comfortable chair and asked to avoid any unnecessary movement or muscle tension. For each block, the two trials containing tempo or pitch changes were excluded from the EEG analyses, thus leaving 10 trials per task and rhythm for subsequent analysis.

Before the EEG session, the parameters for the three tasks were individually adjusted for each participant using a two-down, one-up staircase method, targeting 70.7% accuracy in all tasks (Leek, 2001), separately for the high meter contrast and low meter contrast rhythm. This individual adjustment aimed to make each block equally demanding for the EEG session. These additional trials performed before the EEG session to determine individual parameters also allowed participants to familiarize themselves with the nature of the tasks. For the Pitch and Tempo tasks, the staircase procedure contained a single run where the rhythmic pattern was seamlessly cycled and deviants appeared randomly, separated by at least one intact pattern cycle. Participants were instructed to press a button as soon as they detected a deviant. Button presses within 1 second were considered hits, otherwise the response was considered a miss. The procedure finished after 6 reversals. The threshold was determined as the average deviant magnitude at the last 4 reversals. This procedure was carried out separately for the high and low meter contrast rhythm. For the Visual task, participants were asked to mentally sum 5 sequentially presented numbers with the same parameters as in the EEG session. This was done in discrete trials, and the mean inter-stimulus interval for each trial was adjusted according to the correctness of participant’s response on the previous trial. The procedure finished after 6 reversals (threshold estimated as the average inter-stimulus interval at the last 4 reversals), or after 20 trials (threshold taken as the mean of any available reversals, or the value from the last trial).

After the EEG session, participants rated the subjective difficulty of each task on a discrete scale from 1 (easy) to 7 (difficult).

### EEG recording and preprocessing

The EEG was recorded using a Biosemi Active-Two system (Biosemi, Amsterdam, Netherlands) with 64 Ag-AgCl electrodes placed on the scalp according to the international 10/20 system, and two additional electrodes attached to the mastoids. Head movements were monitored using an accelerometer with two axes (front-back and left-right) attached to the EEG cap and recorded as 2 additional channels. The signals were digitized at 8192-Hz sampling rate, which was high enough to capture distortion-product frequencies relevant for the early auditory responses (Skoe and Kraus, 2010).

#### Analysis of higher-level EEG responses

Higher-level EEG responses refer to EEG activity measured in a low-frequency range (here, at 5 Hz and below), thus corresponding to the frequency range of the actual envelope modulations in the rhythmic inputs (see Figure 2). These responses were analyzed by first downsampling the EEG signals offline to 512 Hz. The continuous EEG signals were then high-pass filtered at 0.1 Hz (4th-order Butterworth filter) to remove slow drifts from the signals. Artifacts related to eye blinks and horizontal eye movements were identified and removed using independent component analysis (Bell and Sejnowski, 1995; Jung et al., 2000) based on visual inspection of their typical waveform shape and topographic distribution. One component was removed for 7 participants, two components for 9 participants, and 9 components for one participant (the eye-movement related activity was clearly distributed across a larger number of components for this participant). Channels containing excessive artifacts or noise were manually selected and linearly interpolated across all trials and conditions, separately for each participant (1 channel for 3 participants, 2 channels for 1 participant). The data were then segmented into 33.6-s long epochs, starting from the onset of the sound sequence in each trial and re-referenced to the average of the 66 channels. The mastoid channels were included because they were expected to prominently capture the responses to auditory rhythms based on previous studies (Nozaradan et al., 2012, 2016b; Lenc et al., 2018, 2020). This was indeed the case, as revealed by the topographical distributions shown in Figure 1. Thus including the mastoid electrodes would enhance the overall signal-to-noise ratio (SNR) of the EEG spectra after averaging across all channels (see below).

**Figure 2.**
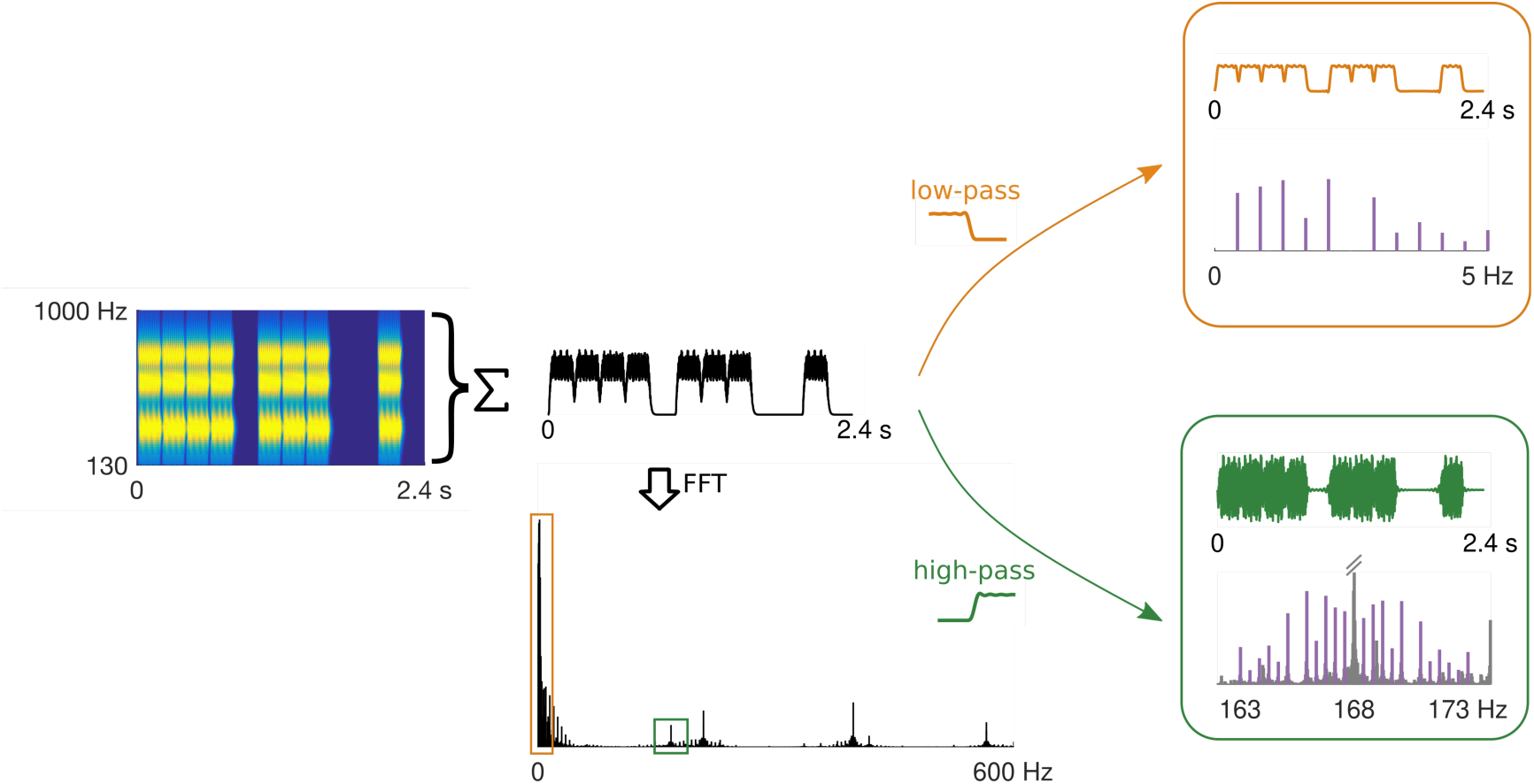
Diagram showing dissociation between higher-level and early auditory EEG responses. Cochleogram on the left shows a response to one cycle of the low meter contrast rhythm across a population of cochlear channels tuned to different frequencies (obtained using the model of Bruce et al., 2018). Summing the peristimulus time histogram across all cochlear channels yields a composite response (shown in black in the middle). The FFT of this composite response (shown on the bottom) reveals how the whole population tracks amplitude envelope modulations (concentrated in the low frequency portion of the spectrum, i.e. at the exact amplitude modulation frequencies), but also phase-locks to the fine structure of the sound input (higher frequency range in the spectrum, at the actual frequencies of the partials and distortion products). Because the fine structure is itself amplitude modulated, the spectrum of the modulator (i.e. amplitude envelope) is reflected in symmetrical sidebands surrounding each partial and distortion product frequency, in line with the shifting theorem of the Fourier Transform (Oppenheim and Schafer, 2009). Thus, the two responses can be separated in the frequency domain by zooming onto the relevant portions of the spectrum, as depicted by the orange rectangle (for the higher-level response) and the green rectangle (for the early auditory response at 168-Hz distortion product). To isolate the higher-level response in the time domain (orange waveform, top right), low-pass filter can be applied to the peristimulus time histogram, which is assumed to take place along the auditory pathway (Chandrasekaran and Kraus, 2010). The early auditory response (green waveform, bottom right) can be isolated in the time domain by high-pass filtering the peristimulus time histogram.

#### Analysis of early auditory EEG responses

Early auditory EEG responses refer to EEG activity measured in a much higher frequency range than the higher-level responses (> 150 Hz), thus corresponding to the frequency range of the actual partials conveying the envelope modulations of the rhythmic inputs (see Figure 2). Here, preprocessing did not include independent component analysis and channel interpolation. The data at the original sampling rate (8192 Hz) were re-referenced to the average of mastoid electrodes, and only signals from three fronto-central channels (Fz, FCz, Cz) were kept for further analyses. Based on previous studies, this standard montage was expected to most strongly capture the auditory frequency-following responses (Skoe and Kraus, 2010; Nozaradan et al., 2016c, 2018).

The preprocessed data were averaged in the time domain across the 10 trials separately for each participant and condition. Time-domain averaging was performed to increase the signal-to-noise ratio by cancelling signals that were not time-locked to the stimulus, while preserving evoked responses elicited by the stimulus, as well as any ongoing activity entrained by the stimulus, which were both assumed to be stationary across trials.

The averaged signals were transformed into the frequency domain using FFT. The obtained spectra were considered to consist of (i) activity elicited by the auditory stimulus, concentrated within narrow peaks and (ii) residual background noise smoothly distributed across a broad range of frequencies (Mouraux et al., 2011; Retter and Rossion, 2016). The contribution of broadband noise was therefore minimized by subtracting the average amplitude at neighboring bins on both sides relative to each frequency bin (bins 2-5 for the higher-level responses and 3-10 for the early auditory responses). A narrower range of bins used for the higher-level responses was to avoid bias in the noise estimate due to prominent 1/f in the lower part of the EEG spectrum. For the early auditory responses, two (instead of one) directly adjacent bins were excluded from the noise estimate due to potential FFT leakage of the response (as the tagged frequencies were not exactly centered on a single frequency bin).

The noise-subtracted spectra were averaged across all channels (66 channels for higher-level responses, 3 channels for early auditory responses) separately for each condition and participant. The magnitudes of higher-level responses were extracted from the bins centered at the 12 frequencies expected based on the stimulus modulation spectrum (see section Frequency-tagging analysis). Magnitudes of the early auditory responses were estimated at the frequencies of the distortion products and their corresponding sideband frequencies (by taking the bin closest to the frequency of interest).

### Overall EEG response magnitude

The overall magnitude of the higher-level responses was estimated as the summed amplitude across all 12 frequencies corresponding to the envelope modulation spectrum of the stimulus (see Figure 3A). The same measure was taken for the early auditory responses by summing across all sideband frequencies, separately for the three distortion-product frequencies (see Figure 4A).

**Figure 3.**
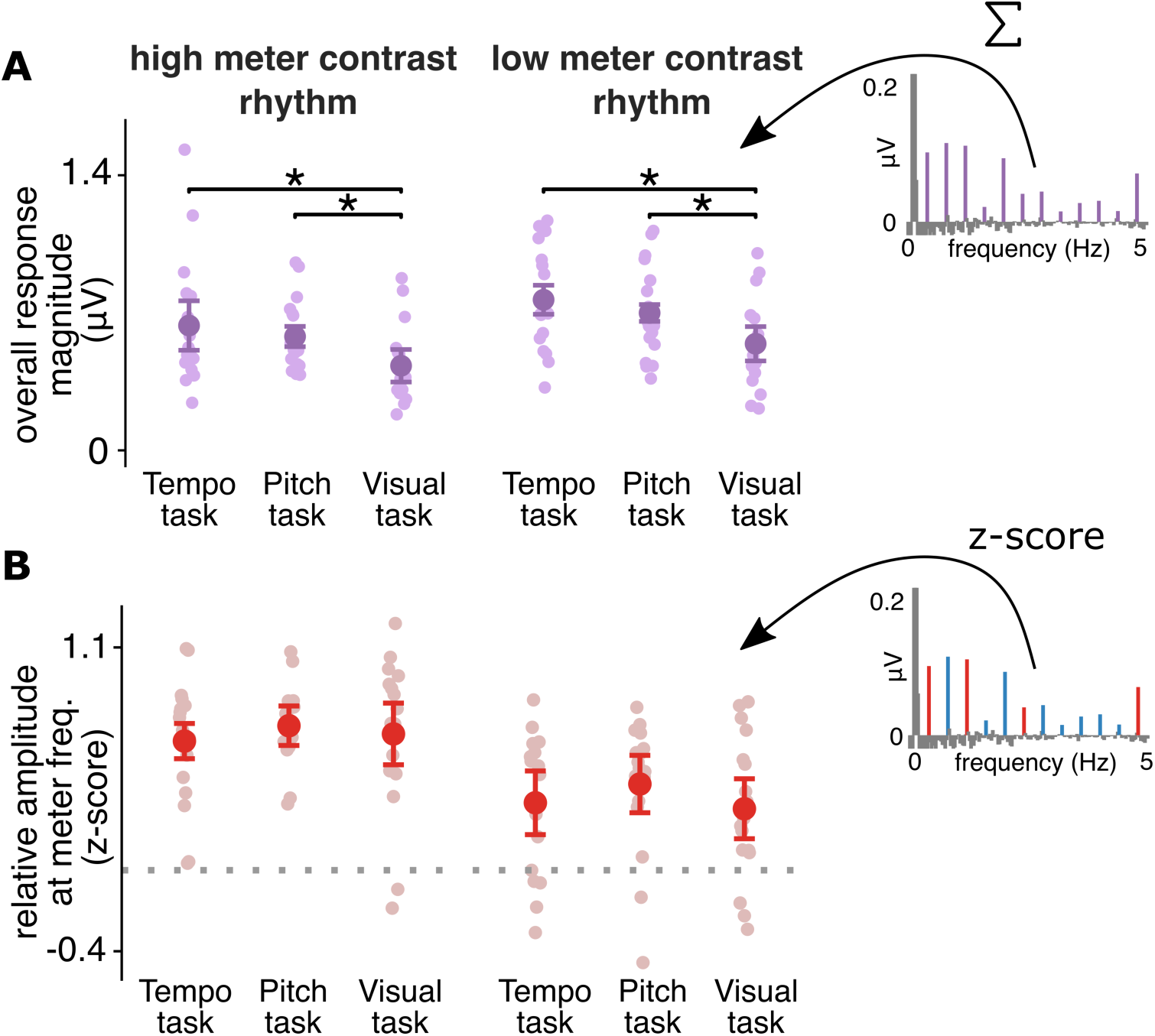
Characteristics of the higher-level EEG responses.The example magnitude spectra on the right visualize how each measure was quantified. Individual participants are shown as lightly shaded data points. Error bars represent 95% CIs (Morey, 2008). (A) Overall response magnitude for the higher-level EEG responses. The amplitudes of the response at all 12 frequencies corresponding to the modulation spectrum of the sound were summed and compared across conditions. For both rhythms, the response was significantly lower during Visual task compared to the two other tasks involving attention to the auditory stimulus (marked by asterisks). (B) Prominence of meter frequencies (mean z-scored amplitude at meter-related frequencies) in the higher-level EEG responses. There was no difference across the three tasks for either rhythm (BF_10_ < 0.3). Moreover, the z-scores show prominent meter frequencies even in response to the rhythm with low contrast at meter frequencies in the acoustic input. The horizontal dashed line represents zero.

**Figure 4.**
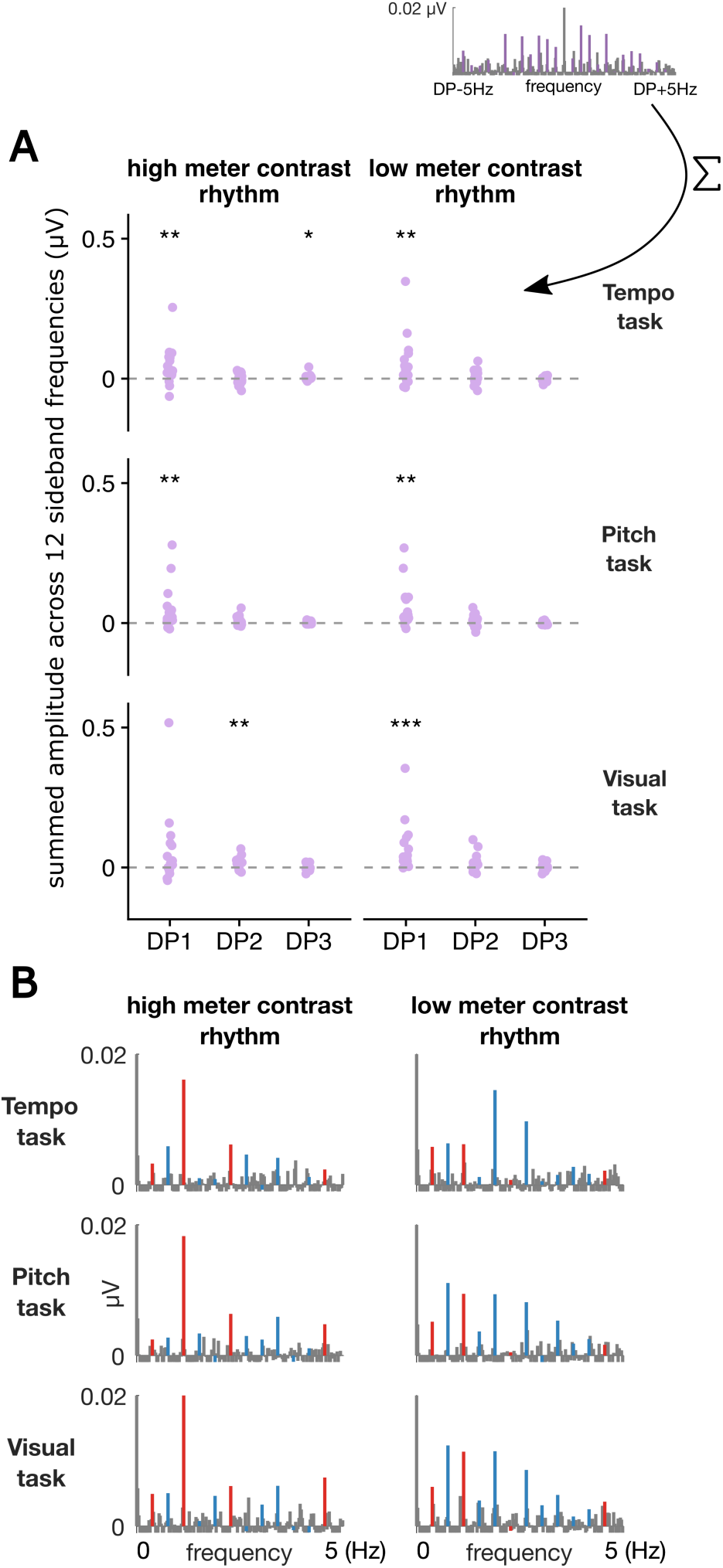
Characteristics of the early auditory EEG responses.(A) Summed early auditory response amplitude averaged across all sidebands, separately for each distortion product frequency (DP1 = 168 Hz, DP2 = 189 Hz, DP3 = 357 Hz). The example magnitude spectrum on the top illustrates how the measure was quantified. Purple data points represent individual participants. Asterisks indicate the statistical significance level of the response when tested against zero (grey dashed line) across participants. * P < 0.05, ** P < 0.01, *** P < 0.001 (Wilcoxon signed rank test, FDR corrected). (B) Spectra of early auditory responses (average across all participants) plotted for the 168 Hz distortion product (DP1) after the corresponding symmetrical sidebands elicited at stimulus modulation frequencies were averaged. Meter-related frequencies are shown in red, meter-unrelated frequencies in blue. The frequency axis is normalized by subtracting the distortion-product frequency for better comparison with the higher-level EEG responses.

To make sure the differences in the overall response magnitude were not due to increased noise floor obscuring the sound-evoked responses, we carried out a control analysis using amplitudes from frequency-bins at positions offset by +7 (i.e. ∼0.21 Hz) relative to the bins centered at the frequencies of interest. These were extracted from the EEG spectra obtained without any noise subtraction, and therefore provided an estimate of the broadband noise level across conditions. This control analysis was only performed for the higher-level responses, as no significant differences in overall response magnitude were found in the early auditory responses (see Results section).

Because the responses at sideband frequencies were generally small, particularly for the sidebands flanking higher distortion-product frequencies, the overall magnitude for the early auditory responses was first compared to zero separately for the sidebands flanking each distortion product to assess whether a significant response was elicited. The validity of this test relies on the fact that, because the spectra were noise-subtracted, an absence of response should result in magnitudes distributed around zero. Only responses at sidebands flanking the lowest distortion product at 168 Hz were consistently above zero (see Table S1 and Figure 4A) across all rhythms and tasks. Therefore, all further analyses of the early auditory responses were carried out only on this distortion-product frequency and corresponding sidebands.

**Table 1.** Comparison of mean z-scored amplitude at meter-related frequencies between higher-level EEG responses and models of auditory subcortical processing using one-sample t-tests.

| EEG variable | rhythm | model | task | mean_model | mean_eeg | sd_eeg | t | df | p |
| --- | --- | --- | --- | --- | --- | --- | --- | --- | --- |
| higher-level response | high meter contrast rhythm | broadband | pitch | 0.81 | 0.71 | 0.19 | -1.99 | 16 | 0.988 |
|  |  |  | tempo | 0.81 | 0.64 | 0.31 | -2.28 | 16 | 0.988 |
|  |  |  | visual | 0.81 | 0.67 | 0.38 | -1.47 | 16 | 0.988 |
|  |  | UREAR_AN | pitch | 0.52 | 0.71 | 0.19 | 4.15 | 16 | 0.0010 *** |
|  |  |  | tempo | 0.52 | 0.64 | 0.31 | 1.61 | 16 | 0.112 |
|  |  |  | visual | 0.52 | 0.67 | 0.38 | 1.69 | 16 | 0.101 |
|  |  | UREAR_IC_BMF2 | pitch | 0.83 | 0.71 | 0.19 | -2.36 | 16 | 0.988 |
|  |  |  | tempo | 0.83 | 0.64 | 0.31 | -2.51 | 16 | 0.988 |
|  |  |  | visual | 0.83 | 0.67 | 0.38 | -1.65 | 16 | 0.988 |
|  |  | UREAR_IC_BMF4 | pitch | 0.79 | 0.71 | 0.19 | -1.58 | 16 | 0.988 |
|  |  |  | tempo | 0.79 | 0.64 | 0.31 | -2.01 | 16 | 0.988 |
|  |  |  | visual | 0.79 | 0.67 | 0.38 | -1.25 | 16 | 0.988 |
|  |  | UREAR_IC_BMF8 | pitch | 0.75 | 0.71 | 0.19 | -0.787 | 16 | 0.988 |
|  |  |  | tempo | 0.75 | 0.64 | 0.31 | -1.51 | 16 | 0.988 |
|  |  |  | visual | 0.75 | 0.67 | 0.38 | -0.846 | 16 | 0.988 |
|  |  | UREAR_IC_BMF16 | pitch | 0.80 | 0.71 | 0.19 | -1.76 | 16 | 0.988 |
|  |  |  | tempo | 0.80 | 0.64 | 0.31 | -2.13 | 16 | 0.988 |
|  |  |  | visual | 0.80 | 0.67 | 0.38 | -1.35 | 16 | 0.988 |
|  |  | UREAR_IC_BMF32 | pitch | 0.81 | 0.71 | 0.19 | -2.09 | 16 | 0.988 |
|  |  |  | tempo | 0.81 | 0.64 | 0.31 | -2.34 | 16 | 0.988 |
|  |  |  | visual | 0.81 | 0.67 | 0.38 | -1.51 | 16 | 0.988 |
|  |  | UREAR_IC_BMF64 | pitch | 0.68 | 0.71 | 0.19 | 0.655 | 16 | 0.447 |
|  |  |  | tempo | 0.68 | 0.64 | 0.31 | -0.601 | 16 | 0.988 |
|  |  |  | visual | 0.68 | 0.67 | 0.38 | -0.105 | 16 | 0.896 |
|  | low meter contrast rhythm | broadband | pitch | 0.06 | 0.43 | 0.33 | 4.66 | 16 | 0.0004 *** |
|  |  |  | tempo | 0.06 | 0.33 | 0.35 | 3.31 | 16 | 0.005 ** |
|  |  |  | visual | 0.06 | 0.30 | 0.35 | 2.88 | 16 | 0.010 * |
|  |  | UREAR_AN | pitch | -0.13 | 0.43 | 0.33 | 6.96 | 16 | <0.0001 *** |
|  |  |  | tempo | -0.13 | 0.33 | 0.35 | 5.48 | 16 | 0.0001 *** |
|  |  |  | visual | -0.13 | 0.30 | 0.35 | 5 | 16 | 0.0003 *** |
|  |  | UREAR_IC_BMF2 | pitch | -0.04 | 0.43 | 0.33 | 5.93 | 16 | <0.0001 *** |
|  |  |  | tempo | -0.04 | 0.33 | 0.35 | 4.5 | 16 | 0.0005 *** |
|  |  |  | visual | -0.04 | 0.30 | 0.35 | 4.04 | 16 | 0.001 ** |
|  |  | UREAR_IC_BMF4 | pitch | -0.29 | 0.43 | 0.33 | 9.01 | 16 | <0.0001 *** |
|  |  |  | tempo | -0.29 | 0.33 | 0.35 | 7.42 | 16 | <0.0001 *** |
|  |  |  | visual | -0.29 | 0.30 | 0.35 | 6.89 | 16 | <0.0001 *** |
|  |  | UREAR_IC_BMF8 | pitch | -0.27 | 0.43 | 0.33 | 8.76 | 16 | <0.0001 *** |
|  |  |  | tempo | -0.27 | 0.33 | 0.35 | 7.18 | 16 | <0.0001 *** |
|  |  |  | visual | -0.27 | 0.30 | 0.35 | 6.66 | 16 | <0.0001 *** |
|  |  | UREAR_IC_BMF16 | pitch | -0.08 | 0.43 | 0.33 | 6.42 | 16 | <0.0001 *** |
|  |  |  | tempo | -0.08 | 0.33 | 0.35 | 4.97 | 16 | 0.0003 *** |
|  |  |  | visual | -0.08 | 0.30 | 0.35 | 4.5 | 16 | 0.0005 *** |
|  |  | UREAR_IC_BMF32 | pitch | 0.05 | 0.43 | 0.33 | 4.73 | 16 | 0.0004 *** |
|  |  |  | tempo | 0.05 | 0.33 | 0.35 | 3.37 | 16 | 0.004 ** |
|  |  |  | visual | 0.05 | 0.30 | 0.35 | 2.94 | 16 | 0.010 ** |
|  |  | UREAR_IC_BMF64 | pitch | 0.01 | 0.43 | 0.33 | 5.28 | 16 | 0.0002 *** |
|  |  |  | tempo | 0.01 | 0.33 | 0.35 | 3.89 | 16 | 0.002 ** |
|  |  |  | visual | 0.01 | 0.30 | 0.35 | 3.45 | 16 | 0.004 ** |

. P < 0.1, \* P < 0.05, \*\* P < 0.01, \*\*\* P < 0.001, FDR corrected

### Relative EEG response at meter frequencies

To assess the relative prominence of specific frequencies in the higher-level responses, amplitudes at the 12 frequencies corresponding to the stimulus modulation spectrum were converted to z-scores, in the same way as for the models of subcortical auditory processing (see section Frequency-tagging analysis). For the early auditory responses, amplitudes corresponding to the same modulation frequency were first averaged across the symmetrical positive and negative sidebands and the resulting 12 values were converted to z-scores (only for 168-Hz distortion-product frequency, see Table S1 and Figure 4A). Higher z-score at a specific frequency indicated that the response at that frequency stood out more prominently relative to the whole set of 12 frequencies elicited by the auditory stimulus.

The z-scores were averaged separately for the meter-related frequencies (frequencies where metric pulses are consistently perceived for these rhythms across listeners, i.e. 0.416, 1.25, 2.5, 5 Hz) and meter-unrelated frequencies (the remaining 8 frequencies in the stimulus modulation spectrum). The mean z-score at meter-related frequencies was taken as a measure of contrast at these frequencies in the neural response, and was compared across conditions.

#### Comparison of EEG with models of subcortical auditory processing

To assess whether the observed EEG responses could be explained by nonlinearities at the early stages of the auditory pathway, the elicited higher-level responses were directly compared to the sound representation estimated by the auditory models. First, the relative prominence of meter frequencies was calculated separately for each model of subcortical auditory processing by taking the mean z-score at meter-related frequencies. Then, the mean meter-related z-scores obtained from the higher-level EEG responses across participants were compared to the corresponding meter z-score from each auditory model, separately for each rhythm and task, using a one-sample t-test. The same analysis was performed for the early auditory responses to confirm that these responses could be largely explained by the auditory models.

#### Comparison of higher-level and early auditory responses

To compare the contrast at meter-related frequencies between early and later processing stages in the same participants, mean z-scored amplitude at these frequencies was compared between the higher-level and early auditory responses across tasks and rhythms.

Moreover, because the highest meter-related frequency (5 Hz) seemed selectively attenuated in the early auditory responses (see Figure 4B), we carried out a control analysis to make sure the differences between higher-level and early auditory responses were not solely driven by differences in the low-pass characteristics of the two responses. The z-scores were re-calculated using only 11 frequencies of interest, i.e. after excluding the amplitude at 5 Hz from the set. Subsequently, only z-scores at 0.416, 1.25, and 2.5 Hz were considered meter-related, and their average was taken to estimate how prominent these frequencies were relative to the whole set of 11 frequencies in the control analysis. This control measure was then used to repeat the comparison between higher-level and early auditory responses, as well as their respective comparisons to the models of subcortical auditory processing.

### Behavioral analyses

Responses to the Tempo and Pitch task from the EEG session were transformed into the sensitivity index d-prime using the equation Z(hit rate) - Z(false alarm rate), where Z is the inverse of the normal cumulative distribution function (Stanislaw and Todorov, 1999). To avoid infinite values, hit rates and false alarm rates with values 1 and 0 were converted to 1/(2N) and 1-1/(2N) respectively (where N is the number of trials on which the proportion is based, Macmillan and Creelman, 2005). The response accuracy on the Visual task was calculated by taking the root-mean-square deviation from the correct response (i.e. the sum of the sequentially-presented number) across trials and averaging across the two rhythm conditions.

### Statistical analyses

The statistical analyses were performed in R (version 3.6.1). Comparisons of EEG measures across conditions were implemented using linear mixed models with lme4 package (version 1.1-21, Bates et al., 2015). The main effects of Rhythm (high meter contrast, low meter contrast) and Task (Tempo, Pitch, Visual), and their interaction were included as fixed effects. For the comparison between the higher-level and early auditory responses, the factor Response (higher-level, early auditory) was also included in the model. Each participant was included as a random-effect intercept. The model comparison was carried out with the *Anova* function from package car (version 3.0-3), using F-tests. Post-hoc comparisons on the fitted models were conducted using emmeans package (version 1.4). Degrees of freedom were approximated using the Kenward-Roger approach and Bonferroni correction was used to adjust the post-hoc test for multiple comparisons. Nonparametric Wilcoxon signed rank tests were used to compare the behavioural responses between conditions, and to assess the significance of the early auditory overall response magnitude against zero (FDR correction for multiple comparisons was used in the latter case). One-sample t-tests were used to compare the meter-related z-scores from the EEG data to the models of subcortical auditory processing (one-tailed, testing EEG > model; p-values were adjusted for multiple comparisons using FDR).

In addition to the null-hypothesis significance tests, we calculated Bayes factors (BF_10_) to express the probability of data under alternative hypothesis (H1) relative to null hypothesis (H0), as implemented in BayesFactor package (version 0.9.12-4.2). We considered a BF_10_ > 3 as evidence in favour of the alternative hypothesis and BF_10_ < 0.3 as evidence in favour of the null hypothesis (Jeffreys, 1998; Lee and Wagenmakers, 2014).

## Results

### Behavioral results

Participants successfully detected the deviants during the EEG session for both the Pitch task (mean d’ [SD] for the high meter contrast rhythm = 1.23 [0.92], for the low meter contrast rhythm = 1.28 [0.55]), and the Tempo task (mean d’ [SD] for the high meter contrast rhythm = 1.28 [0.77], for the low meter contrast rhythm = 1.28 [1.10]). There was no difference in the sensitivity between the Pitch and Tempo task (Ps > 0.25, BFs_10_ < 0.6). In the Visual task, the root-mean-square error of the responses averaged across rhythms was 87.1 (SD = 71.6), suggesting that the task was difficult and participants might have often missed some numbers in the sequence. This was in line with participants rating the Visual task as much more difficult than the Pitch task (Wilcoxon signed rank test, two-sided P = 0.0007), or the Tempo task (Wilcoxon signed rank test, P = 0.0003). One participant’s data were excluded from the calculation of the error, due to the misunderstanding of the instructions - instead of summing, the participant was concatenating the presented numbers and memorizing the sequence. However, the EEG data of this participant were not excluded, as she reported similar difficulty for the Visual task as the rest of the participants.

### Overall EEG response magnitude

Figure 1 shows the spectra of the higher-level EEG responses elicited by the rhythmic stimuli across the three tasks. As shown in Figure 3A, the summed amplitude of the higher-level responses across all twelve frequencies was significantly different across the three tasks (F_2,80_ = 16.3, P < 0.0001, BF_10_ > 100). This was due to smaller overall amplitude in the Visual task relative to the Tempo task (β = −0.21, t_82_ = −5.61, P < 0.0001, 95% CI = [-0.31, −0.12]), and the Pitch task (β = −0.15, t_82_ = −4, P = 0.0004, 95% CI = [-0.25, −0.06]). The corresponding analysis performed on the shifted frequency bins where no signal was expected indicated that broadband noise amplitude was comparable across conditions (no significant main effect of Task: P = 0.62, BF_10_ = 0.13, and no interaction Task x Rhythm: P = 0.39, BF_10_ = 0.32). Thus, increased noise alone could not account for the observed overall response decrease in the Visual task.

Figure 4B shows the spectra of the early auditory EEG responses. Unlike for the higher-level responses, the summed amplitude across all sidebands for the early auditory responses did not differ across conditions (Ps > 0.22, BFs_10_ < 0.21). This suggested that attentional focus did not affect the putatively earlier stage of sound processing, and only emerged at later stages.

### Relative EEG response at meter frequencies

The relative prominence of meter frequencies in the elicited higher-level EEG responses was significantly larger for the high than low meter contrast rhythm (F_1,80_ = 38.9, P < 0.0001, BF_10_ > 100), as expected based on the physical structure of the rhythmic stimuli (see Methods section). However, as shown in Figure 3B, z-scores at meter frequencies were comparable across tasks (no significant main effect of Task: P = 0.32, BF_10_ = 0.24, and no interaction Task x Rhythm: P = 0.79, BF_10_ = 0.18). Similarly, meter-related frequencies were significantly more prominent in the early auditory responses to the high meter contrast rhythm (F_1,80_ = 42.8, P < 0.0001, BF_10_ > 100), but there was no effect of task (Ps > 0.65, BFs_10_ < 0.13).

### Comparison of EEG responses with models of subcortical auditory processing

For the high meter contrast rhythm, the relative prominence of meter-related frequencies in the higher-level EEG responses was explained by all considered models of subcortical auditory processing (see Figure 5 and Table 1). The mean z-score at meter-related frequencies measured in the elicited EEG was not significantly different from either model, except there was a significantly greater z-score for the EEG responses in the pitch task when compared to the UR_EAR model of the auditory nerve response. However, for the low meter contrast rhythm, meter-related frequencies were consistently more prominent in the elicited EEG responses than predicted by all models of subcortical auditory processing. Importantly, this was the case even when participants were carrying out the visual task. These results were further corroborated by a control analysis showing that meter-related frequencies in the higher-level responses were robustly enhanced even when the highest meter frequency (5 Hz) was excluded from the analysis (see Table S2).

**Figure 5.**
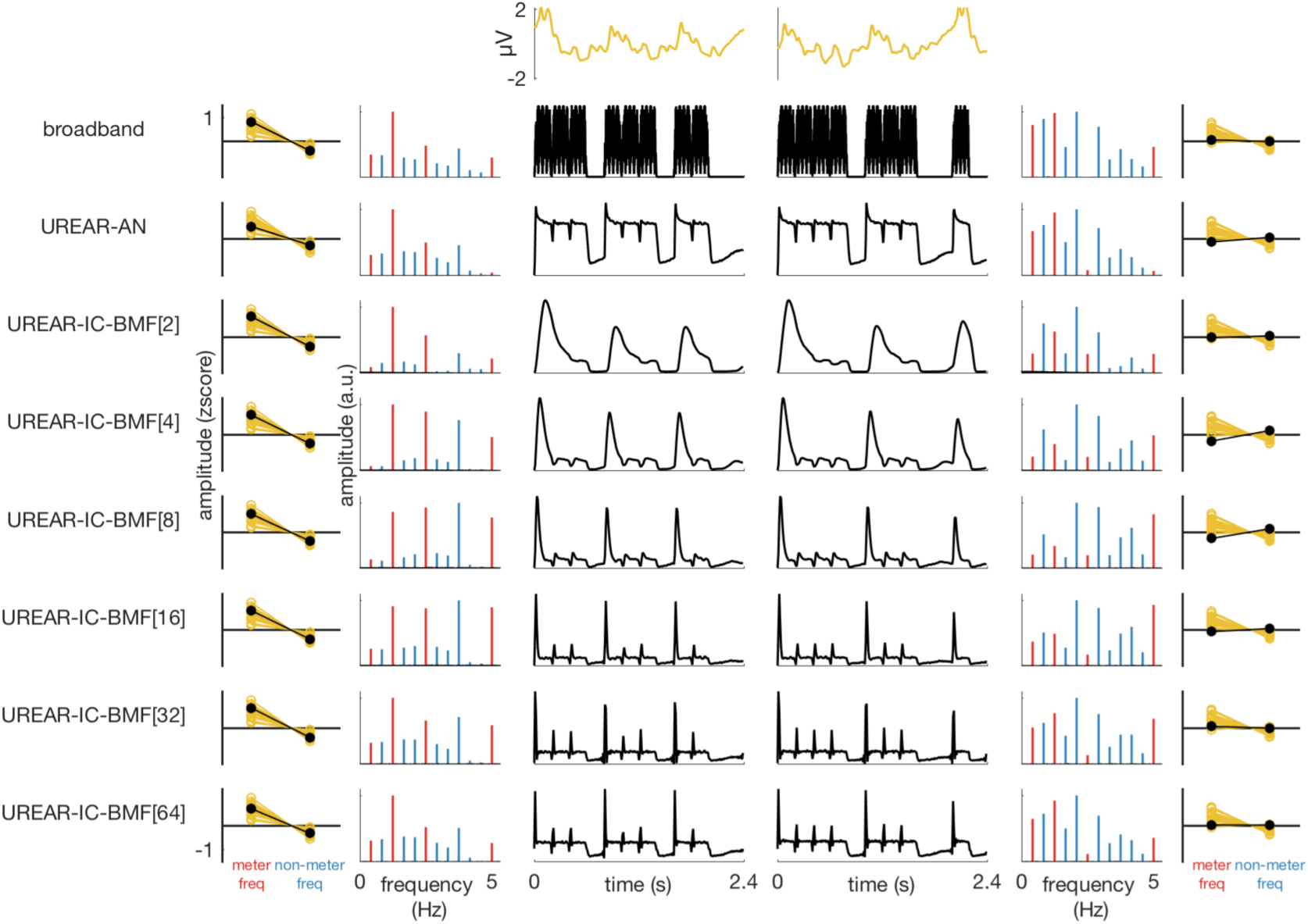
Comparison of higher-level EEG responses with models of auditory subcortical processing. Data for the high meter contrast rhythms are shown on the left, and data for the low meter contrast rhythm are shown on the right. The higher-level EEG response to one pattern cycle averaged across all pattern repetitions, trials, tasks, and participants is depicted on the top in yellow. This response was extracted after low-pass filtering at 30 Hz, and averaging 9 frontocentral channels (F1, F2, Fz, C1, C2, Cz, FC1, FC2, FCz). Each row below the EEG response corresponds to the output of one model of subcortical auditory processing. The model labels are shown on the left (depending on the parameter settings, the IC cell simulated with UR_EAR had a specific best modulation frequency, which is listed in square brackets in Hz). In the center of the figure are the responses of the models to one cycle of the rhythmic pattern depicted as mean firing rate across time. The mean rate over time was transformed into the frequency domain using FFT, and the resulting spectra are shown next to the time-domain responses. Meter-related frequencies are shown in red, and meter-unrelated frequencies in blue. On the sides of the figure are the z-scored spectral amplitudes averaged separately across meter-related and meter-unrelated frequencies. The yellow data points represent EEG responses of individual participants (z-scores averaged across the 3 tasks), and the black data points represent the auditory model. While the z-scores at meter-related frequencies did not differ reliably between the models and the EEG responses for the high meter contrast rhythm on the left, the EEG response at these frequencies was selectively enhanced for the low meter contrast rhythm (right).

As expected, the same set of comparisons for the early auditory responses showed no clear differences from the models of subcortical auditory processing (see Table S3). Even though the meter frequencies in the early auditory responses were significantly above some auditory models after excluding the highest meter-related frequency in the control analysis (UR-EAR-IC with best modulation frequencies 4, 8, 16 Hz, see Table S4), this was not systematic (in contrast to the higher-level responses), and was related to a selective suppression of the lower meter frequencies in these models compared to the rest of the models (see Figure 5).

Together, these results indicate that, as expected, the contrast at meter frequencies in the EEG response to the rhythmic input with prominent meter frequencies in its acoustic structure was mainly driven by low-level processing of this input. However, these early processing stages cannot fully explain the EEG response to the rhythmic input with less prominent meter frequencies in its acoustic structure. Most importantly, the processes responsible for the selective enhancement of meter frequencies for this latter rhythm seemed to be involved to a similar degree across the three attentional tasks.

### Comparison of higher-level and early auditory EEG responses

A mixed model with factors Rhythm, Task and Response revealed a main effect of Rhythm (F_1,176_ = 68.03, P < 0.0001, BF_10_ > 100), as expected from the separate analyses of the higher-level and early auditory responses above, suggesting that both types of response were sensitive to the physical structure of the auditory input. There was also a main effect of Response (F_1,176_ = 58.5, P < 0.0001, BF_10_ > 100). As shown in Figure 6, the relative prominence of meter frequencies was consistently larger in the higher-level responses across all tasks and both rhythms. Moreover, this could not be easily explained by differences in the low-pass characteristic of the responses, as the same results were obtained in a control analysis where the highest meter-related frequency (i.e. 5 Hz) was excluded (see Supplementary Material). This suggests that there was a significant enhancement of meter-related frequencies at the later processing stage indexed by the higher-level responses.

**Figure 6.**
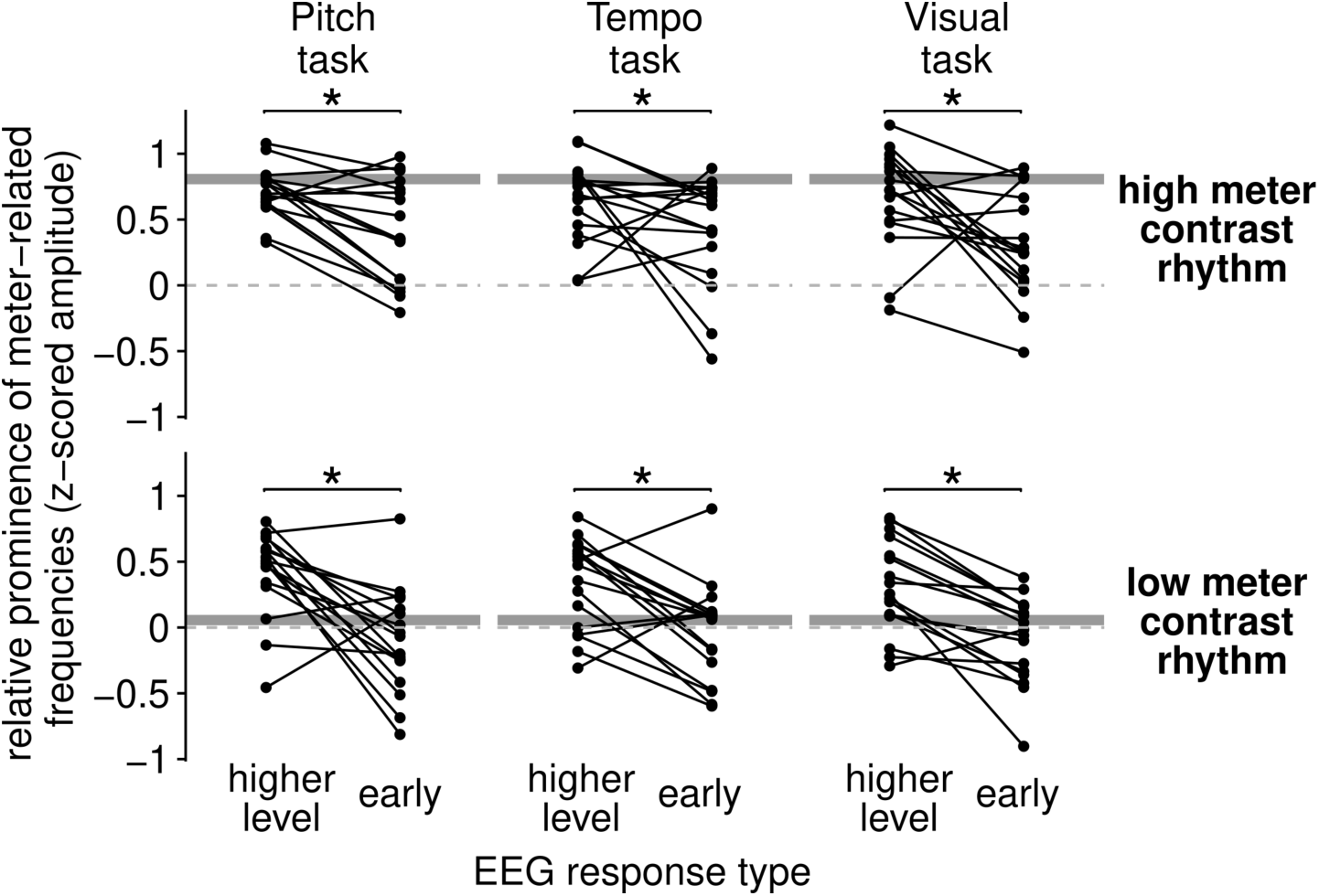
Comparison of prominence of meter-related frequencies in the higher-level and early auditory EEG responses.The mean z-scored amplitude at meter-related frequencies is plotted separately for each task, rhythm, and EEG response type. Individual data points represent participants. Horizontal continuous grey lines correspond to the mean z-scores at meter frequencies taken from the broadband amplitude envelope of the corresponding acoustic stimulus. The horizontal dashed grey lines represent zero (i.e. equal relative prominence of meter-related and meter-unrelated frequencies). The meter-related frequencies were consistently more prominent in the higher-level EEG responses across all rhythms and task (main effect of response type, indicated by asterisks).

## Discussion

Our results show that while attentional focus affects the overall sensitivity of the brain to auditory rhythmic inputs, it has little influence on the selective contrast at meter periodicities in the elicited neural response. Moreover, while the magnitude of this selective contrast in the EEG response was readily explained by low-level auditory processing for the rhythm with prominent meter frequencies in the acoustic input, this was not the case for the rhythm that lacked prominent contrast at meter periodicities in its physical structure. Together, these results suggest the critical engagement of high-level processes that shape the neural representation of a rhythmic input by selectively enhancing contrast at meter periodicities across behavioral contexts even in rhythms where this contrast is not prominent. These results add to the evidence that rhythm perception is shaped by a range of processes including higher-level cortical stages, with different degrees of flexibility and automaticity.

### Wide range of low-level and higher-level processes in meter perception

A number of studies have consistently shown that brain can selectively enhance meter-related frequencies, particularly for low meter contrast rhythms. However, the behavioral context varied across these studies. Some asked participants to report small occasional changes in the duration of a sound event making up the rhythm (Nozaradan et al., 2012, 2017b; Lenc et al., 2018). While those small changes in duration of single time intervals are difficult to detect at a local scale, their detection is easier on a global scale, as the deviant interval results in a misalignment of the subsequent sequence with the perceived meter. Therefore, this task implicitly encourages participants to rely on an internal metric structure (Schulze, 1978; Jones and Yee, 1997; Grube and Griffiths, 2009). Similarly, some studies asked participants to focus on the tempo (overall perceived speed) of the sequences, either searching for local transient changes (Lenc et al., 2020), of for later comparison with a test sequence (Tal et al., 2017). Other studies have asked participants to report any temporal irregularities in the stimuli while none were actually present (Nozaradan et al., 2018), or to simply attend to the sequences (Nozaradan et al., 2016a). Our results suggest that selective neural enhancement of meter frequencies can take place even without attention directly focused on the temporal properties of the stimulus (as observed during the pitch task). Interestingly, this enhancement was also observed when sound was ignored altogether, i.e., even when the overall signal-to-noise ratio (SNR) of the response was decreased. Nevertheless, our results inform future frequency-tagging studies, which may find it advantageous to employ behavioral tasks that encourage participants to focus on the auditory stimuli, even when the task-relevant dimension is orthogonal to rhythm processing, to increase the SNR and facilitate estimation of response properties.

While our results suggest that neural responses to rhythmic input might involve processes largely independent on attentional focus, this does not imply that neural processing of rhythm is fixed and inflexible. A number of recent studies have shown that selective neural enhancement of meter periodicities reflects flexible processes, which are sensitive to mental imagery (Nozaradan et al., 2011; Li et al., 2019), non-temporal features of the acoustic input (Lenc et al., 2018), prior experience (Chemin et al., 2014), and recent context (Lenc et al., 2020). Importantly, our current findings show that the internal transformation of a rhythmic input towards a particular metric category can be flexibly enhanced, possibly changed, but it is difficult to suppress completely: it has an automatic component. Whether the robust component can be changed by long-term exposure remains to be seen in future studies (Hannon et al., 2012b, 2012a; London et al., 2017; van der Weij et al., 2017; Polak et al., 2018).

These findings thus reveal similarities between meter processing and other higher-level auditory processes, such as auditory stream segregation or change detection (Sussman, 2017). While automatic in some contexts (Woods et al., 1992; Alho et al., 1994; Dyson et al., 2005; Teki et al., 2011, 2016; Masutomi et al., 2016), attention can boost or bias these processes (Haroush et al., 2010; Auksztulewicz and Friston, 2015; O’Sullivan et al., 2015; Costa-Faidella et al., 2017), particularly when the input provides ambiguous sensory cues to the system (Sussman et al., 2007; Gutschalk et al., 2015). Similarly, the brain response to isochronous auditory sequences might be spontaneously shaped by the intrinsic preference for binary structures (Brochard et al., 2003; Pablos Martin et al., 2007), but this can be biased towards different forms of organization by top-down processes dependent on attentional resources (Nozaradan et al., 2011; Chemin et al., 2014; Celma-Miralles and Toro, 2019). Importantly, some higher-level auditory processes can be largely suppressed (especially by high load) but not completely abolished (Woldorff et al., 1991; Alain and Izenberg, 2003; Chait et al., 2012; Billig and Carlyon, 2016; Molloy et al., 2019). Together, these studies suggest that when assessing effects of attention on a particular perceptual phenomenon, it is important to keep in mind that (i) multiple processes can be involved (Chait et al., 2012), (ii) these different processes may be differentially affected by different kinds of tasks (Bidet-Caulet et al., 2007; Yerkes et al., 2019), and (iii) the effect of task may also depend on the sensory information provided by the stimulus (Gutschalk et al., 2015). With respect to meter perception, identification of the underlying internal processes and the types of resources they rely on remains worthwhile goal for future investigations (Lenc et al., 2018).

### Dissociation between overall gain and selective contrast at meter periodicities

An overall decrease is sensitivity to sound input while carrying out a visual task has been previously reported by a large number of studies measuring early event-related potentials (Woods et al., 1992; Alho et al., 1994; Okamoto et al., 2011), frequency tagged responses (Keitel et al., 2011, 2013; Riecke et al., 2014), or BOLD activations in sensory cortices (Petkov et al., 2004; Shomstein and Yantis, 2004; Johnson and Zatorre, 2005; Riecke et al., 2017). In line with these studies, we observed increased overall gain of the higher-level EEG responses during the auditory tasks in comparison to the visual task. In other words, there was a general increase in the amplitude of neural activity time-locked to the rhythmic auditory stimulus. However, this non-specific enhancement could simply represent a proportional increase of magnitude across the response spectrum, thus being equivalent to a multiplicative enhancement of the response in the time domain. In that case, the relative contrast in the response at meter periodicities should necessarily remain constant.

As opposed to such non-specific enhanced gain of the whole response, a change in the selective contrast at meter periodicities would demonstrate selective increase at meter-related frequencies. The important distinction between overall gain and selective contrast at meter frequencies has been often neglected in studies claiming to measure “neural entrainment to meter” (e.g. Tierney and Kraus, 2014; Hickey et al., 2020). Our results provide a cautionary example, whereby non-specific increase in response to the sound trivially explained by attention could have been misinterpreted as an enhancement of meter periodicities in brain responses. Instead, our method allowed us to dissociate between these two accounts, exploiting the fact that energy in the modulation spectra of our complex rhythmic stimuli is not solely distributed across meter-related frequencies. For this reason, using non-isochronous rhythms, particularly rhythms with less energy at meter frequencies, is advantageous over strictly isochronous stimuli where changes in overall response gain and selective contrast enhancement cannot be differentiated. In addition, using z-score normalization instead of the difference in raw spectral magnitude makes the measure robust to multiplicative gain (which would yield greatest raw magnitude increase at frequencies already prominent in the spectra).

### Robust responses at meter periodicities even with low meter contrast in the input

Despite the significant differences in the gain of the higher-level EEG responses, the selective contrast at meter frequencies was not affected by task. However, this finding could be trivially explained by passive matching of stimulus modulation structure in the neural response. While faithful tracking of stimulus envelope is fundamental for auditory perception (Peelle et al., 2013; Di Liberto et al., 2018; Etard and Reichenbach, 2019; Ghinst et al., 2019), the brain must go beyond one-to-one representation of the sensory input to achieve adaptive behavior (Kuchibhotla and Bathellier, 2018). Thus, the sensory input is continuously transformed within the brain towards higher-level categories (Ley et al., 2014; Brodbeck et al., 2018; Rossion et al., 2020; Sankaran et al., 2020; Yin et al., 2020). Such transformations are critical for timing perception already at the level of single intervals (Desain and Honing, 2003; Jacoby and McDermott, 2017), but also patterns of intervals (Notter et al., 2018), and for meter perception, where a range of physically different acoustic inputs can be mapped onto the same set of periodic pulses (Nozaradan et al., 2017a). To assess whether the internal processes involved in this transformation were engaged even when attention was withdrawn from the auditory input, we compared the higher-level EEG responses to simulated representation of the auditory input across different auditory subcortical stages. While keeping in mind that absence of evidence is not evidence of absence, our results suggests that well-described low-level nonlinearities in early auditory pathway cannot fully account for the cortical brain response to the low meter contrast rhythm.

This was further confirmed by the early sensory responses measured in the same EEG as the high-level responses. Even though the tagged frequencies used to identify the early sensory responses in the current study were most likely not in a high enough frequency range to strictly isolate brainstem responses from cortical responses (Coffey et al., 2016; Holmes et al., 2018), these responses may have preferentially captured contributions from primary auditory fields, as well as subcortical nuclei (Chandrasekaran and Kraus, 2010; Nourski and Brugge, 2011). Therefore, comparing EEG responses tagged at low frequencies (higher-level responses) vs. high frequencies (early auditory responses) may still be a useful way to separate sound representation in early auditory cortices from responses originating in a wide network of structures involved in rhythm processing (Patel and Iversen, 2014; Merchant et al., 2015).

Our results are in line with Nozaradan et al. (2018) who observed a similar enhancement of meter-related frequencies in the higher-level EEG responses compared to the early auditory EEG responses. While they only observed higher-level response enhancement for the low meter contrast rhythm, this was the case for both rhythms in the current study. However, if our current results were related to low SNR for early auditory responses resulting in attenuation of the most prominent peaks in the spectra, one would expect to find opposite effects on meter-related frequencies for the two rhythms due to the differences in their physical structure. Moreover, non-selective attenuation of high frequencies in the early auditory responses alone did not fully explain the smaller prominence of meter frequencies when compared to the higher-level EEG responses. Finally, the prominence of meter frequencies in the early auditory responses was strongly modulated by the type of rhythm, showing sensitivity to the acoustic structure of the input. Therefore, together with the output of subcortical auditory processing models, our results provide evidence that (i) higher-level processes further enhance contrast at meter frequencies, especially when these meter frequencies are not prominent in the auditory input, and (ii) these processes remain active even when overall responsiveness to sound input is decreased (e.g. during a demanding visual task).

Recently, it has been proposed that low-level nonlinearities such as adaptation, amplitude-modulation tuning, and heightened sensitivity to contrast in early stages of the auditory pathway could predict whether and what metric structure is perceived in a rhythmic stimulus (Rajendran et al., 2017; Zuk et al., 2018), and the consistency of perceived meter across individual listeners (Rajendran et al., 2020). While these low-level phenomena are definitely important in shaping rhythm perception, such models are inherently limited to enhancement of sensitivity to contrast that is *already present* in the physical input. Indeed, these models are based on the long-standing assumption in psychology and neuroscience (likely driven by over-emphasis on Western classical and popular music) that meter perception is driven by temporal contrasts defined by acoustic properties of the sound input (Longuet-Higgins and Lee, 1984; Povel and Essens, 1985; Jones and Boltz, 1989; Palmer and Krumhansl, 1990; Drake et al., 2000; Toiviainen and Snyder, 2003). Strong arguments against these assumptions have been recently raised by a number of authors (see, e.g. London et al., 2017; van der Weij et al., 2017). Indeed, such models will unlikely explain perception of musical genres where the phase (e.g. reggae, ska, swing, mazurka) or period (e.g. tresillo, cascara, or rumba clave in afro-cuban music) of the perceived metric structure is weakly cued in the temporal distribution of features in the physical sound. Instead, over-constrained views may in the end lose explanatory power by ignoring diversity and flexibility in the cognitive phenomenon across cultures in pursuit of a reductionist mechanistic explanation.

The weak explanatory power of biologically plausible models of subcortical auditory processing to account for our EEG results adds to the evidence that meter perception involves higher-level transformations of the input, providing flexibility within (Repp, 2007; Repp et al., 2008; Chemin et al., 2014; Lenc et al., 2020) and across individuals (McKinney and Moelants, 2006; Martens, 2011; Hannon et al., 2012a; Kalender et al., 2013; Polak et al., 2018; Witek et al., 2020). Instead of offering a mechanistic explanation for the current EEG results, we emphasize the need for more data and powerful designs, in order to thoroughly describe the perceptual phenomenon in question, and how it is shaped by input features, behavioral goals, context, exposure, and learning. Similarly, we do not claim that the contrast at meter frequencies measured in EEG responses is one-to-one with meter perception in a phenomenological sense. At the same time, it is important to note that all measures of perception are indirect (including behavioral measures), and critically depend on the definition of the perceptual phenomenon (see also Rossion et al., 2020). If meter is defined as the perception of pulses that are time-locked to the temporal structure of the stimulus, and if pulse is understood as something that consistently occurs at regularly-spaced time points *and not otherwise* (thus creating a temporal contrast), our EEG measure is directly relevant for meter processing. Moreover, our analysis was directly informed by tapping data from previous studies, thus constraining the set of behaviorally relevant periodicities based on an additional measure of meter perception.

The fact that we observed significantly enhanced responses at meter frequencies in the low meter contrast rhythm irrespective of attentional focus may seem inconsistent with the fMRI study of Chapin et al. (Chapin et al., 2010). In that study, participants listened to rhythms that had few acoustic cues to meter periodicities but still elicited stable meter perception. The authors observed larger BOLD responses within a network of structures typically associated with meter perception when participants were actively listening (memorizing the rhythm for subsequent reproduction) than when they were memorizing a visual array of letters. However, this result could potentially be explained by non-specific changes in the BOLD response due to auditory stimulation interacting with attention, or task-related motor preparation. Based on the current results, we suggest that the processing of low meter contrast rhythms may not be inherently different from high meter contrast rhythms. Indeed, rhythms with little acoustic cues to meter are ubiquitous across cultures (Cohn, 2016; London et al., 2017; Witek, 2017; Câmara and Danielsen, 2018). Hence, such rhythmic inputs may help to reveal the transformations that take place within the brain when the sensory input is mapped onto an internal metric representation, and eventually behavioral output, while controlling for acoustic or low-level confounds.

### Evidence for robust meter processing complementary to MMN studies of passive listening

Our observation that processes related to meter perception are engaged robustly across behavioral contexts is consistent with previous studies using the mismatch-negativity event-related potential, or MMN (Ladinig et al., 2009; Winkler et al., 2009; Bouwer et al., 2014, 2016). Complementary to these studies, we observed task-independent neural enhancement of meter frequencies even for the low meter contrast rhythm, while MMN responses have only been assessed for rhythms with very prominent acoustic cues to meter periodicities. Because MMN paradigms rely on assumptions about predictions and regularity violations, which are not well-defined for meter perception (see e.g. London et al., 2017),

MMN studies involve uncertainty about the type of deviation that may elicit differential responses depending on its timing relative to the perceived pulse (Bouwer and Honing, 2015). MMN studies therefore rely on statistically linking the perceived pulse with salient sound events, and hence limit themselves to study of high meter contrast rhythms. Moreover, typical MMN studies employ passive listening with low cognitive load and no control of participant’s attentional focus (Sussman et al., 2014), whereas we directly manipulated the attentional state of the listener with active demanding tasks that significantly affected the overall magnitude of the EEG responses and also parieto-occipital alpha power (see Supplementary Material), which is an established index of crossmodal attentional engagement (Fu et al., 2001; Jensen and Mazaheri, 2010; Mo et al., 2011; Mazaheri et al., 2014). Our results thus represent an important step towards describing whether processes involved in meter perception depend on limited resources, which may be shared across modalities and cognitive domains (Marois and Ivanoff, 2005; Chait et al., 2012; Murphy et al., 2017; Molloy et al., 2020).

### Conclusions

The human auditory system possesses a remarkable capacity to carry out high-level processes with limited attentional resources (Murphy et al., 2017). The current study provides evidence that this may also be the case for processes involved in meter perception in the context of musical rhythm. Our results indicate that the brain selectively emphasizes perceptually relevant periodicities even when overall sensitivity to sound is decreased due to a distracting task. Moreover, such neural emphasis occurs when the periodicities are not prominent in the sensory input, and their enhancement is not readily accounted for by low-level auditory processing. Therefore, these robust neural processes may support the spontaneity of meter perception when listening to a variety of musical inputs, while still allowing for flexibility and context dependence in meter perception within and across individuals.

## Supplementary material

### Control analysis of higher-level and early auditory EEG responses excluding the highest (5 Hz) frequency

To make sure that the differences in the prominence of meter-related frequencies between higher-level and early auditory EEG responses were not solely driven by low-pass biases, we re-calculated the relative prominence of meter frequencies after excluding the highest frequency of interest (5 Hz) from the set. This frequency corresponded to rate of individual events in the rhythms, but captured also harmonics of the slower perceived metric pulses. Yet, the amplitude at this frequency would be affected most prominently if higher frequencies in the response were broadly attenuated irrespective of their contribution to the contrast at meter periodicities (e.g. within a neural network behaving like a simple low-pass filter). A mixed model with factors Rhythm, Task and Response still revealed a main effect of Rhythm (F_1,176_ = 53.4, P < 0.0001, BF_10_ > 100) and Response (F_1,176_ = 35.0, P < 0.0001, BF_10_ > 100), thus replicating the results from the main analysis. This indicates that the enhanced selective contrast at meter periodicities in the higher-level responses cannot be fully explained by non-selective enhancement of higher frequencies (i.e. without considering their relevance for the perceived meter).

**Table S1.**
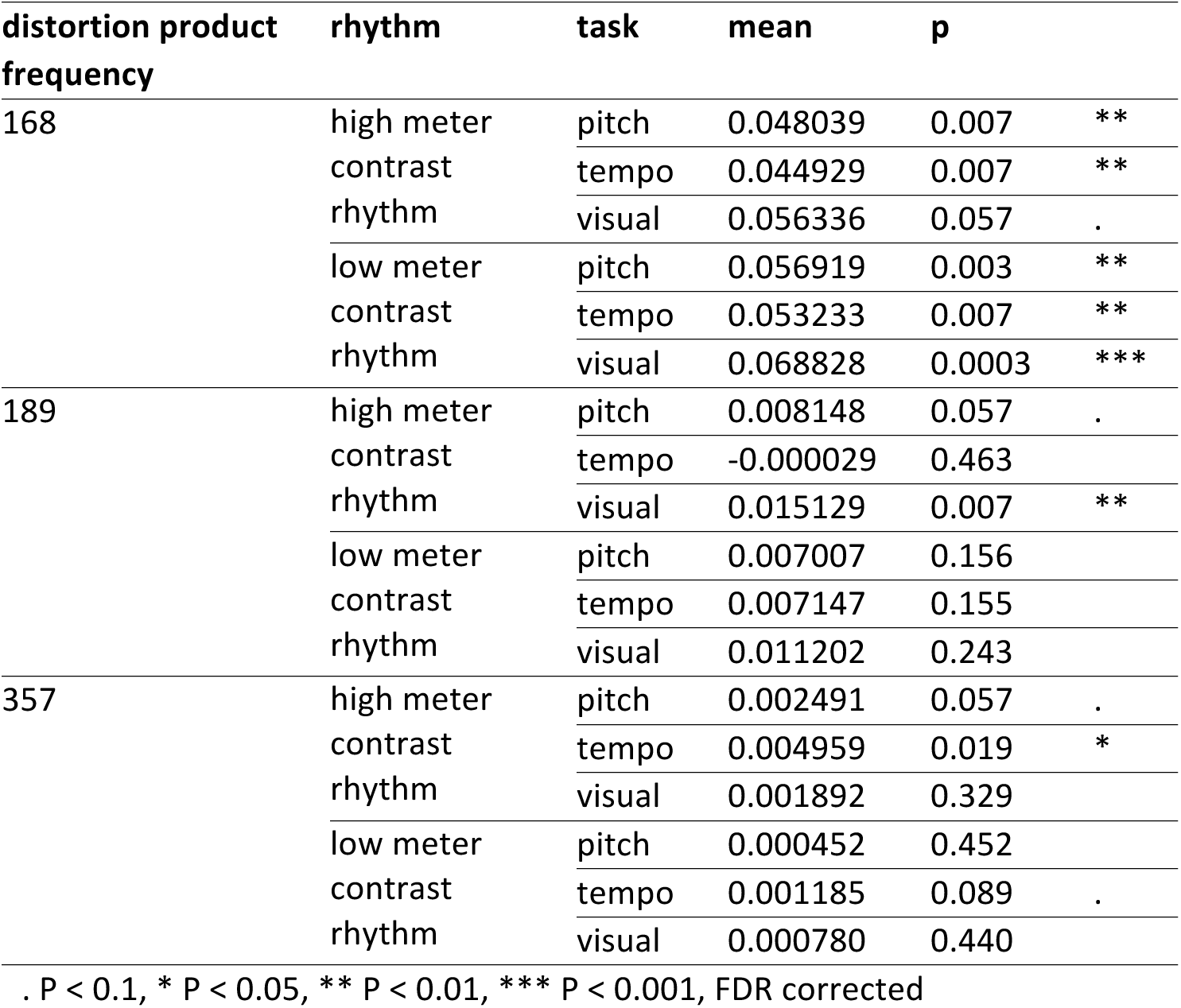
Comparison of mean early auditory response amplitude averaged across all sidebands against zero using non-parametric Wilcoxon signed rank tests.

**Table S2.** Control analysis of mean z-scored amplitude at meter-related frequencies without taking the highest frequency (5 Hz) into account. Comparison between higher-level EEG responses and models of auditory subcortical processing using one-sample t-tests.

| EEG variable | rhythm | model | task | mean_model | mean_eeg | sd_eeg | t | df | p |
| --- | --- | --- | --- | --- | --- | --- | --- | --- | --- |
| higher-level response | high meter contrast rhythm | broadband | pitch | 1.06 | 0.84 | 0.23 | -4.03 | 16 | 1.000 |
|  |  |  | tempo | 1.06 | 0.78 | 0.39 | -2.98 | 16 | 1.000 |
|  |  |  | visual | 1.06 | 0.76 | 0.48 | -2.63 | 16 | 1.000 |
|  |  | UREAR_AN | pitch | 0.97 | 0.84 | 0.23 | -2.27 | 16 | 1.000 |
|  |  |  | tempo | 0.97 | 0.78 | 0.39 | -1.96 | 16 | 1.000 |
|  |  |  | visual | 0.97 | 0.76 | 0.48 | -1.79 | 16 | 1.000 |
|  |  | UREAR_IC_BMF2 | pitch | 1.06 | 0.84 | 0.23 | -4.02 | 16 | 1.000 |
|  |  |  | tempo | 1.06 | 0.78 | 0.39 | -2.97 | 16 | 1.000 |
|  |  |  | visual | 1.06 | 0.76 | 0.48 | -2.62 | 16 | 1.000 |
|  |  | UREAR_IC_BMF4 | pitch | 0.90 | 0.84 | 0.23 | -1.08 | 16 | 1.000 |
|  |  |  | tempo | 0.90 | 0.78 | 0.39 | -1.28 | 16 | 1.000 |
|  |  |  | visual | 0.90 | 0.76 | 0.48 | -1.23 | 16 | 1.000 |
|  |  | UREAR_IC_BMF8 | pitch | 0.76 | 0.84 | 0.23 | 1.48 | 16 | 0.171 |
|  |  |  | tempo | 0.76 | 0.78 | 0.39 | 0.20 | 16 | 0.748 |
|  |  |  | visual | 0.76 | 0.76 | 0.48 | -0.01 | 16 | 0.865 |
|  |  | UREAR_IC_BMF16 | pitch | 0.79 | 0.84 | 0.23 | 0.94 | 16 | 0.338 |
|  |  |  | tempo | 0.79 | 0.78 | 0.39 | -0.11 | 16 | 0.899 |
|  |  |  | visual | 0.79 | 0.76 | 0.48 | -0.27 | 16 | 0.967 |
|  |  | UREAR_IC_BMF32 | pitch | 0.91 | 0.84 | 0.23 | -1.20 | 16 | 1.000 |
|  |  |  | tempo | 0.91 | 0.78 | 0.39 | -1.35 | 16 | 1.000 |
|  |  |  | visual | 0.91 | 0.76 | 0.48 | -1.29 | 16 | 1.000 |
|  |  | UREAR_IC_BMF64 | pitch | 0.94 | 0.84 | 0.23 | -1.80 | 16 | 1.000 |
|  |  |  | tempo | 0.94 | 0.78 | 0.39 | -1.69 | 16 | 1.000 |
|  |  |  | visual | 0.94 | 0.76 | 0.48 | -1.57 | 16 | 1.000 |
|  | low meter contrast rhythm | broadband | pitch | 0.13 | 0.51 | 0.40 | 4.01 | 16 | 0.002 ** |
|  |  |  | tempo | 0.13 | 0.40 | 0.39 | 2.85 | 16 | 0.015 * |
|  |  |  | visual | 0.13 | 0.25 | 0.48 | 1.05 | 16 | 0.317 |
|  |  | UREAR_AN | pitch | 0.14 | 0.51 | 0.40 | 3.86 | 16 | 0.002 ** |
|  |  |  | tempo | 0.14 | 0.40 | 0.39 | 2.70 | 16 | 0.019 * |
|  |  |  | visual | 0.14 | 0.25 | 0.48 | 0.93 | 16 | 0.338 |
|  |  | UREAR_IC_BMF2 | pitch | 0.02 | 0.51 | 0.40 | 5.11 | 16 | 0.0002 *** |
|  |  |  | tempo | 0.02 | 0.40 | 0.39 | 3.95 | 16 | 0.002 ** |
|  |  |  | visual | 0.02 | 0.25 | 0.48 | 1.96 | 16 | 0.077 . |
|  |  | UREAR_IC_BMF4 | pitch | -0.46 | 0.51 | 0.40 | 10.16 | 16 | <0.0001 *** |
|  |  |  | tempo | -0.46 | 0.40 | 0.39 | 9.03 | 16 | <0.0001 *** |
|  |  |  | visual | -0.46 | 0.25 | 0.48 | 6.14 | 16 | <0.0001 *** |
|  |  | UREAR_IC_BMF8 | pitch | -0.65 | 0.51 | 0.40 | 12.05 | 16 | <0.0001 *** |
|  |  |  | tempo | -0.65 | 0.40 | 0.39 | 10.94 | 16 | <0.0001 *** |
|  |  |  | visual | -0.65 | 0.25 | 0.48 | 7.71 | 16 | <0.0001 *** |
|  |  | UREAR_IC_BMF16 | pitch | -0.49 | 0.51 | 0.40 | 10.46 | 16 | <0.0001 *** |
|  |  |  | tempo | -0.49 | 0.40 | 0.39 | 9.33 | 16 | <0.0001 *** |
|  |  |  | visual | -0.49 | 0.25 | 0.48 | 6.38 | 16 | <0.0001 *** |
|  |  | UREAR_IC_BMF32 | pitch | -0.08 | 0.51 | 0.40 | 6.18 | 16 | <0.0001 *** |
|  |  |  | tempo | -0.08 | 0.40 | 0.39 | 5.04 | 16 | 0.0002 *** |
|  |  |  | visual | -0.08 | 0.25 | 0.48 | 2.85 | 16 | 0.015 * |
|  |  | UREAR_IC_BMF64 | pitch | 0.13 | 0.51 | 0.40 | 3.98 | 16 | 0.002 ** |
|  |  |  | tempo | 0.13 | 0.40 | 0.39 | 2.82 | 16 | 0.015 * |
|  |  |  | visual | 0.13 | 0.25 | 0.48 | 1.03 | 16 | 0.317 |

. P < 0.1, \* P < 0.05, \*\* P < 0.01, \*\*\* P < 0.001, FDR corrected

**Table S3.** Comparison of mean z-scored amplitude at meter-related frequencies between early auditory EEG responses and models of auditory subcortical processing using one-sample t-tests.

| EEG variable | rhythm | model | task | mean_model | mean_eeg | sd_eeg | t | df | p |
| --- | --- | --- | --- | --- | --- | --- | --- | --- | --- |
| early auditory response | high meter contrast rhythm | broadband | pitch | 0.81 | 0.41 | 0.39 | -4.20 | 16 | 1.0 |
|  |  |  | tempo | 0.81 | 0.42 | 0.42 | -3.80 | 16 | 1.0 |
|  |  |  | visual | 0.81 | 0.32 | 0.41 | -5.00 | 16 | 1.0 |
|  |  | UREAR_AN | pitch | 0.52 | 0.41 | 0.39 | -1.15 | 16 | 1.0 |
|  |  |  | tempo | 0.52 | 0.42 | 0.42 | -0.93 | 16 | 1.0 |
|  |  |  | visual | 0.52 | 0.32 | 0.41 | -2.04 | 16 | 1.0 |
|  |  | UREAR_IC_BMF2 | pitch | 0.83 | 0.41 | 0.39 | -4.38 | 16 | 1.0 |
|  |  |  | tempo | 0.83 | 0.42 | 0.42 | -3.98 | 16 | 1.0 |
|  |  |  | visual | 0.83 | 0.32 | 0.41 | -5.17 | 16 | 1.0 |
|  |  | UREAR_IC_BMF4 | pitch | 0.79 | 0.41 | 0.39 | -4.00 | 16 | 1.0 |
|  |  |  | tempo | 0.79 | 0.42 | 0.42 | -3.61 | 16 | 1.0 |
|  |  |  | visual | 0.79 | 0.32 | 0.41 | -4.80 | 16 | 1.0 |
|  |  | UREAR_IC_BMF8 | pitch | 0.75 | 0.41 | 0.39 | -3.60 | 16 | 1.0 |
|  |  |  | tempo | 0.75 | 0.42 | 0.42 | -3.24 | 16 | 1.0 |
|  |  |  | visual | 0.75 | 0.32 | 0.41 | -4.42 | 16 | 1.0 |
|  |  | UREAR_IC_BMF16 | pitch | 0.80 | 0.41 | 0.39 | -4.09 | 16 | 1.0 |
|  |  |  | tempo | 0.80 | 0.42 | 0.42 | -3.69 | 16 | 1.0 |
|  |  |  | visual | 0.80 | 0.32 | 0.41 | -4.89 | 16 | 1.0 |
|  |  | UREAR_IC_BMF32 | pitch | 0.81 | 0.41 | 0.39 | -4.25 | 16 | 1.0 |
|  |  |  | tempo | 0.81 | 0.42 | 0.42 | -3.85 | 16 | 1.0 |
|  |  |  | visual | 0.81 | 0.32 | 0.41 | -5.04 | 16 | 1.0 |
|  |  | UREAR_IC_BMF64 | pitch | 0.68 | 0.41 | 0.39 | -2.89 | 16 | 1.0 |
|  |  |  | tempo | 0.68 | 0.42 | 0.42 | -2.56 | 16 | 1.0 |
|  |  |  | visual | 0.68 | 0.32 | 0.41 | -3.73 | 16 | 1.0 |
|  | low meter contrast rhythm | broadband | pitch | 0.06 | -0.10 | 0.40 | -1.58 | 16 | 1.0 |
|  |  |  | tempo | 0.06 | -0.04 | 0.38 | -1.03 | 16 | 1.0 |
|  |  |  | visual | 0.06 | -0.09 | 0.33 | -1.84 | 16 | 1.0 |
|  |  | UREAR_AN | pitch | -0.13 | -0.10 | 0.40 | 0.31 | 16 | 1.0 |
|  |  |  | tempo | -0.13 | -0.04 | 0.38 | 0.95 | 16 | 1.0 |
|  |  |  | visual | -0.13 | -0.09 | 0.33 | 0.45 | 16 | 1.0 |
|  |  | UREAR_IC_BMF2 | pitch | -0.04 | -0.10 | 0.40 | -0.54 | 16 | 1.0 |
|  |  |  | tempo | -0.04 | -0.04 | 0.38 | 0.06 | 16 | 1.0 |
|  |  |  | visual | -0.04 | -0.09 | 0.33 | -0.58 | 16 | 1.0 |
|  |  | UREAR_IC_BMF4 | pitch | -0.29 | -0.10 | 0.40 | 2.00 | 16 | 0.3 |
|  |  |  | tempo | -0.29 | -0.04 | 0.38 | 2.72 | 16 | 0.2 |
|  |  |  | visual | -0.29 | -0.09 | 0.33 | 2.49 | 16 | 0.2 |
|  |  | UREAR_IC_BMF8 | pitch | -0.27 | -0.10 | 0.40 | 1.79 | 16 | 0.4 |
|  |  |  | tempo | -0.27 | -0.04 | 0.38 | 2.50 | 16 | 0.2 |
|  |  |  | visual | -0.27 | -0.09 | 0.33 | 2.24 | 16 | 0.2 |
|  |  | UREAR_IC_BMF16 | pitch | -0.08 | -0.10 | 0.40 | -0.13 | 16 | 1.0 |
|  |  |  | tempo | -0.08 | -0.04 | 0.38 | 0.48 | 16 | 1.0 |
|  |  |  | visual | -0.08 | -0.09 | 0.33 | -0.09 | 16 | 1.0 |
|  |  | UREAR_IC_BMF32 | pitch | 0.05 | -0.10 | 0.40 | -1.53 | 16 | 1.0 |
|  |  |  | tempo | 0.05 | -0.04 | 0.38 | -0.98 | 16 | 1.0 |
|  |  |  | visual | 0.05 | -0.09 | 0.33 | -1.77 | 16 | 1.0 |
|  |  | UREAR_IC_BMF64 | pitch | 0.01 | -0.10 | 0.40 | -1.07 | 16 | 1.0 |
|  |  |  | tempo | 0.01 | -0.04 | 0.38 | -0.50 | 16 | 1.0 |
|  |  |  | visual | 0.01 | -0.09 | 0.33 | -1.22 | 16 | 1.0 |

. P < 0.1, \* P < 0.05, \*\* P < 0.01, \*\*\* P < 0.001, FDR corrected

**Table S4.**
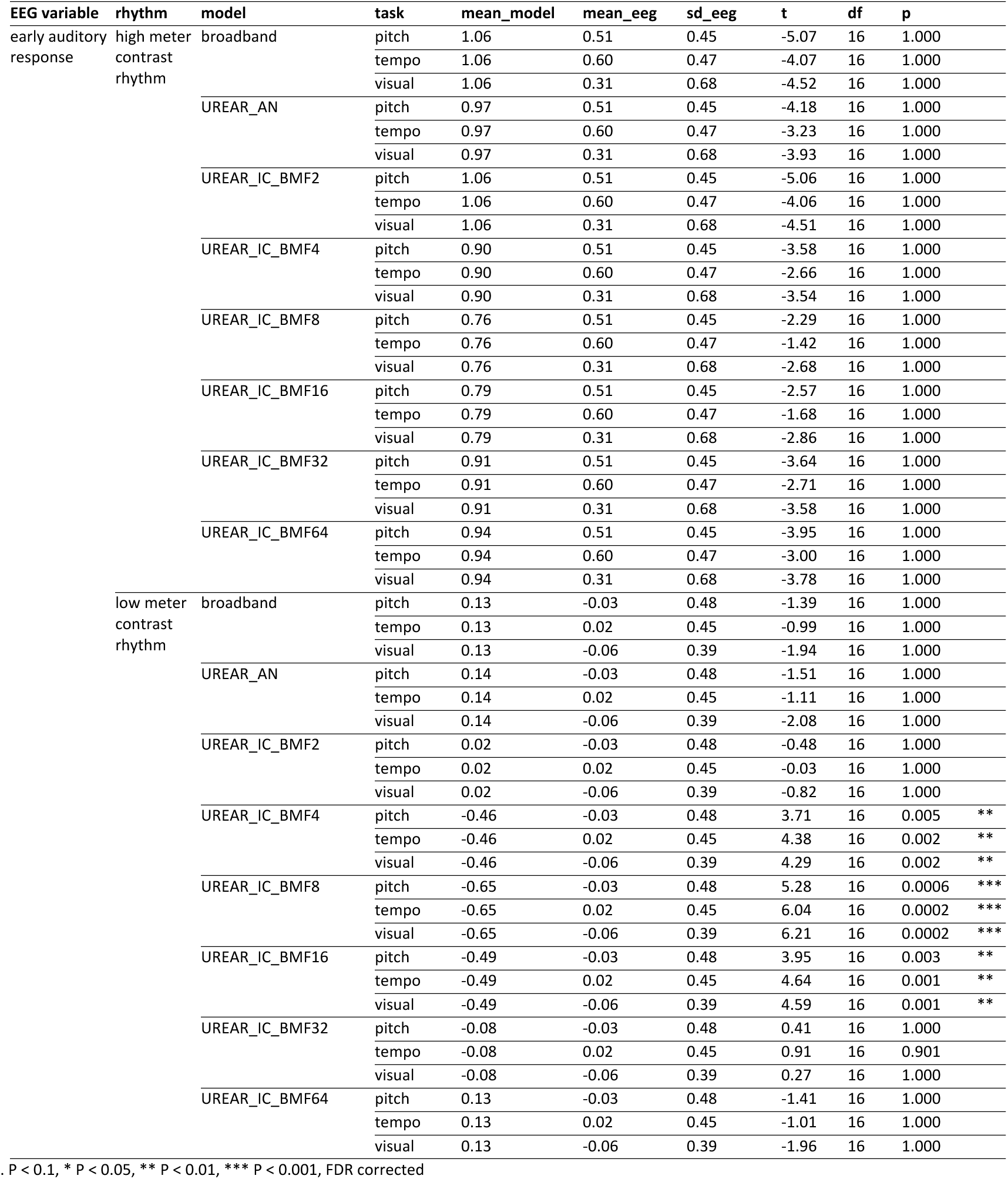
Control analysis of mean z-scored amplitude at meter-related frequencies without taking the highest frequency (5 Hz) into account. Comparison between early auditory EEG responses and models of auditory subcortical processing using one-sample t-tests.

### EEG oscillatory alpha activity

Parieto-occipital alpha power was measured as an independent index of attentional engagement during the visual task, compared to the two auditory tasks. To separate the oscillatory activity from the 1/f background, the time-domain preprocessed data were subjected to irregular-resampling auto-spectral analysis (IRASA) (Wen and Liu, 2016). This method utilizes the fact that irregular resampling with non-integer factors results in shifts of the oscillatory component along the frequency axis, whereas the 1/f component remains constant. The procedure was carried out separately for each channel, condition, and participant. The data from each trial were segmented into 15 overlapping windows that were equally spaced throughout the trial. The number of samples in each window corresponded to the largest power of 2 that did not exceed 90% of trial duration. For each window, the auto-power spectrum was estimated using FFT after multiplication with a Hann function. This was performed for the original sampling rate, and also after resampling using pairs of resampling factors f and 1/f (where f was taken from 0.1 to 0.9 in steps of 0.05). The geometric mean of the auto-power spectra was taken across each pair of resampling factors. The power spectrum of the fractal component was estimated as the median-average spectrum across all values of f, separately for each window. The power spectrum of the fractal component was then averaged across the 15 windows and subtracted from the average power spectrum of the original signal (without any resampling), to obtain an estimate of the oscillatory component.

In the resulting oscillatory power spectra, all frequencies that were expected to contain neural activity elicited by the acoustic stimulus (i.e. harmonics of the pattern repetition rate, 0.416 Hz) were set to zero. Subsequently, the power in the alpha range was quantified by taking the mean power between 8 and 12 Hz (Iemi et al., 2017; van Diepen and Mazaheri, 2017; Van Diepen et al., 2019), separately for each condition. Alpha power was averaged across 18 parieto-occipital channels (Iz, O1, Oz, O2, PO7, PO3, POz, PO4, PO8, P9, P7, P3, P1, Pz, P2, P4, P8, P10) that were expected to show largest effects of cross-modal attention based on previous studies (Fu et al., 2001; Mazaheri et al., 2014; van Diepen and Mazaheri, 2017).

The power of alpha oscillatory activity from the parieto-occipital electrodes was significantly modulated by task (F_2,80_ = 10, P = 0.0001, BF_10_ > 100). As shown in Figure S1, the power significantly decreased during the Visual task compared to the Tempo task (β = −0.03, t_82_ = −4.39, P = 0.0001, 95% CI = [-0.05, −0.01]) and Pitch task (β = - 0.02, t_82_ = −3.08, P = 0.008, 95% CI = [-0.04, −0.005]).

Visual inspection of the data suggested that the effect might have been only present for participants who had higher baseline alpha power. This was supported by a significant improvement of the model after adding the interaction between Task and alpha power in the Visual task as a continuous predictor (F_3,72.5_ = 11.72, P < 0.0001). Across participants, higher power in the Visual-task condition was related to greater power increase in the Tempo task (β = 1.41, t_45.3_ = 5.41, P < 0.0001, 95% CI = [0.73, 1.98]) and in the Pitch task (β = 1.16, t_45.3_ = 4.47, P < 0.0001, 95% CI = [0.49, 1.73]).

**Figure S1.**
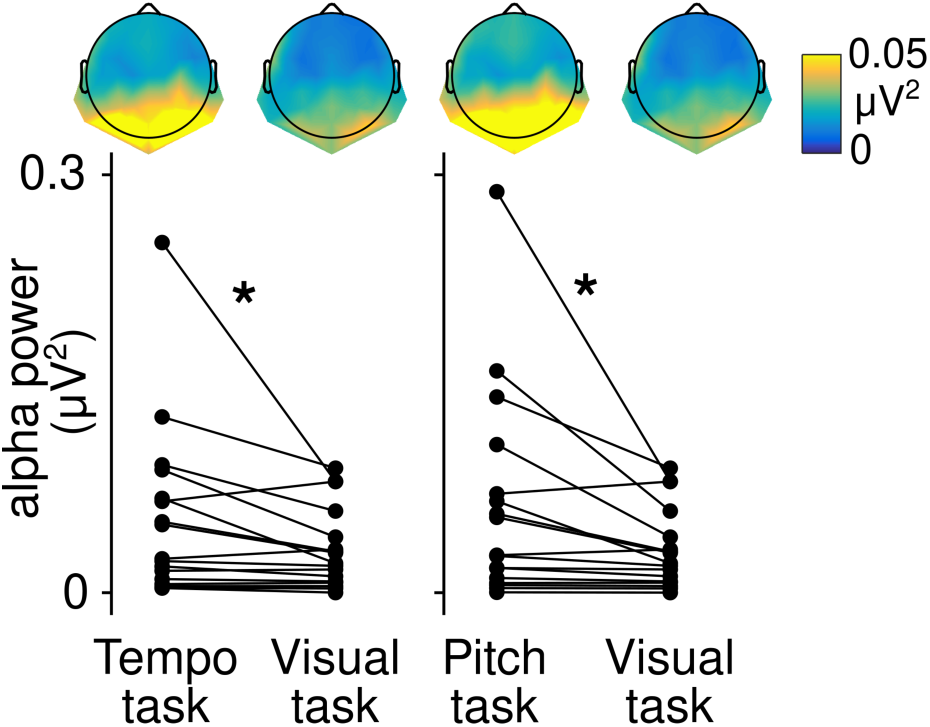
EEG power at the alpha frequency elicited across the different tasks at posterior channels. Alpha power was significantly smaller (p < 0.01, marked by asterisks) during the visual task, and the magnitude of the effect depended on the baseline alpha response across participants.

